# Epigenetic changes induced by developmental PFAS exposure in zebrafish associate with behavioral alterations in unexposed offspring

**DOI:** 10.64898/2026.04.14.718393

**Authors:** Adeolu Z. Ogunleye, Michela Di Criscio, Manon Fallet, Jonas Zetzsche, Coralie Yon, Nikolai Scherbak, Steffen H. Keiter, Philipp Antczak, Joëlle Rüegg

**Author notes:** These authors contributed equally to this work (Adeolu Z. Ogunleye, Michela Di Criscio, Manon Fallet). These authors contributed equally to this work (Steffen H. Keiter, Philipp Antczak, Joëlle Rüegg).

## Abstract

Per- and polyfluoroalkyl substances (PFAS) are widespread environmental contaminants with documented toxic effects, yet their multi- and transgenerational impacts on neurodevelopment and underlying mechanisms remain poorly understood. Here, we present a comprehensive study delineating the effects of developmental exposure to environmentally relevant concentrations of PFOS and PFBS on behavior, transcriptome, and genome-wide DNA methylation patterns in the directly exposed generation (F0) and their unexposed offspring (F1 and F2) in zebrafish. Both PFOS and PFBS altered larval behavior, linked to transcriptomic and DNA methylation changes in neuro-related pathways, even in the unexposed offspring. Importantly, specific DNA methylation changes in F0 were associated with behavioral outcomes in F2 animals, suggesting that these alterations could underlie transgenerational effects. Pathways associated with differentially methylated genes were prominently enriched for response to light and circadian regulation. Our findings demonstrate that developmental exposure to PFAS causes transgenerational behavioral effects in zebrafish and suggest that epigenetic changes induced by direct exposure may serve as markers for predicting outcomes in subsequent, unexposed generations.

**TEASER:** PFAS induce circadian-related epigenetic changes in zebrafish associated with behavioral impacts in unexposed offspring.

## INTRODUCTION

Biological effects resulting from chemical exposure may extend beyond the directly exposed generation, affecting phenotypes and potentially leading to adverse outcomes in unexposed offspring. These effects may manifest as intergenerational inheritance, where germ cells are directly exposed, or as transgenerational inheritance, where no exposure to the chemical occurs. However, our understanding and prediction of these effects remain limited, as most toxicological studies focus primarily on immediate impacts within a single generation.

Epigenetic modifications are hypothesized to mediate inter- and transgenerational phenotypic effects in response to different stimuli, including chemical exposure (*1*–*6*). The terms epigenetic marks and mechanisms refer to structural modifications or molecular processes that affect chromosomal regions. Thereby, these modifications and processes register or maintain changes in transcriptional activity states without involving mutations (*7*). Epigenetic alterations can be inherited across mitosis and meiosis, influencing cell differentiation, development, and ultimately determining the overall transcriptomic and phenotypic profile of an organism (*8*).

Per- and polyfluoroalkyl substances (PFAS) constitute a large class of chemicals known for their toxicological effects (*9*, *10*). The PFAS class encompasses over 12,000 compounds, characterised by a CnF2n+1– structure, known for their resistance to degradation due to the strong carbon-fluorine bond. Since the 1950s, PFAS have been widely used in stain-proof, greaseproof, water-repellent, and fire-resistant products (*11*).

Within this class, Perfluorooctane Sulfonate (PFOS) is one of the most recognized and prevalent PFAS contaminants, found ubiquitously in soil and surface waters (*12*–*14*). Epidemiological studies have found associations between PFOS exposure and various health issues, including immune and hormonal disruptions (*10*), low birth weight (*15*), and altered neurodevelopment (*16*). Animal studies support these findings, showing effects on immunity in mice (*17*) and medaka fish (*18*), thyroid hormone disruption in monkeys (*19*), rats (*20*), and zebrafish (*21*), as well as behavioral changes in zebrafish and mice (*22*–*24*).

Regulatory measures have reduced exposure to PFOS in Europe and the U.S. (*25*, *26*), but have also led to the increased use of alternative compounds with largely unknown long-term toxicity. One such compound, PFBS, has been detected in aquatic ecosystems (0.01–4,520 ng/L) and linked to behavioral changes in zebrafish larvae and adults (*27*).

Previous studies have reported neuro-phenotypical and transcriptional inter- and transgenerational alterations following short-term exposure to PFOS in zebrafish (*23*, *28*), as well as multi-generational thyroidal disruption due to PFBS exposure in medaka (*29*). Moreover, altered epigenetic processes have been linked to direct PFAS exposure (*30*, *31*). However, whether PFAS-induced epigenetic modifications mediate inter- and transgenerational phenotypic effects remains untested. Furthermore, while behavioral alterations have been observed, the molecular mechanisms underlying their inheritance are poorly understood, limiting our ability to predict adverse outcomes in unexposed generations. In this study, we examined whether environmentally relevant concentrations of PFOS and PFBS can induce behavioral, transcriptomic, and epigenomic changes across three generations, covering both inter- and transgenerational inheritance. Additionally, we evaluated the relationship between phenotypic and molecular alterations to explore the potential of molecular changes as predictors of adverse phenotypic outcomes in subsequent, unexposed generations.

## RESULTS

### Developmental exposure to PFAS induces multi- and transgenerational behavioral changes

To assess whether PFOS and PFBS in environmentally relevant concentrations induce inter- and transgenerational effects, we exposed zebrafish to PFOS and PFBS at a lower concentration (LC) of 0.2 µg/L and a higher concentration (HC) of 2 µg/L during the first 28 days of development. Subsequently, we assessed behavioral alterations in directly exposed (F0), germline exposed (F1), and unexposed (F2) larvae using the Larval Photo Motor Response (LPMR) test. Larvae exhibited increased movement in the dark, confirming the validity of the protocols used and indicating that none of the exposure groups showed substantial alterations in general larval response patterns (fig. S1)

Results for all generations and exposures are shown in Fig. 1, A-D. In F0, we observed no difference in the acclimatization phase (LON1) and the recovery phase (LON2) during the LPMR test. Yet, during the stressful phase (LOFF), the movement of the F0 larvae decreased in almost all conditions compared to control (PFOS LC, PFOS HC, and PFBS LC: adjusted p-value < 0.0001), apart from exposure to PFBS HC, which induced an increase in larvae movement (adjusted p-value = 0.005).

**Fig. 1.**
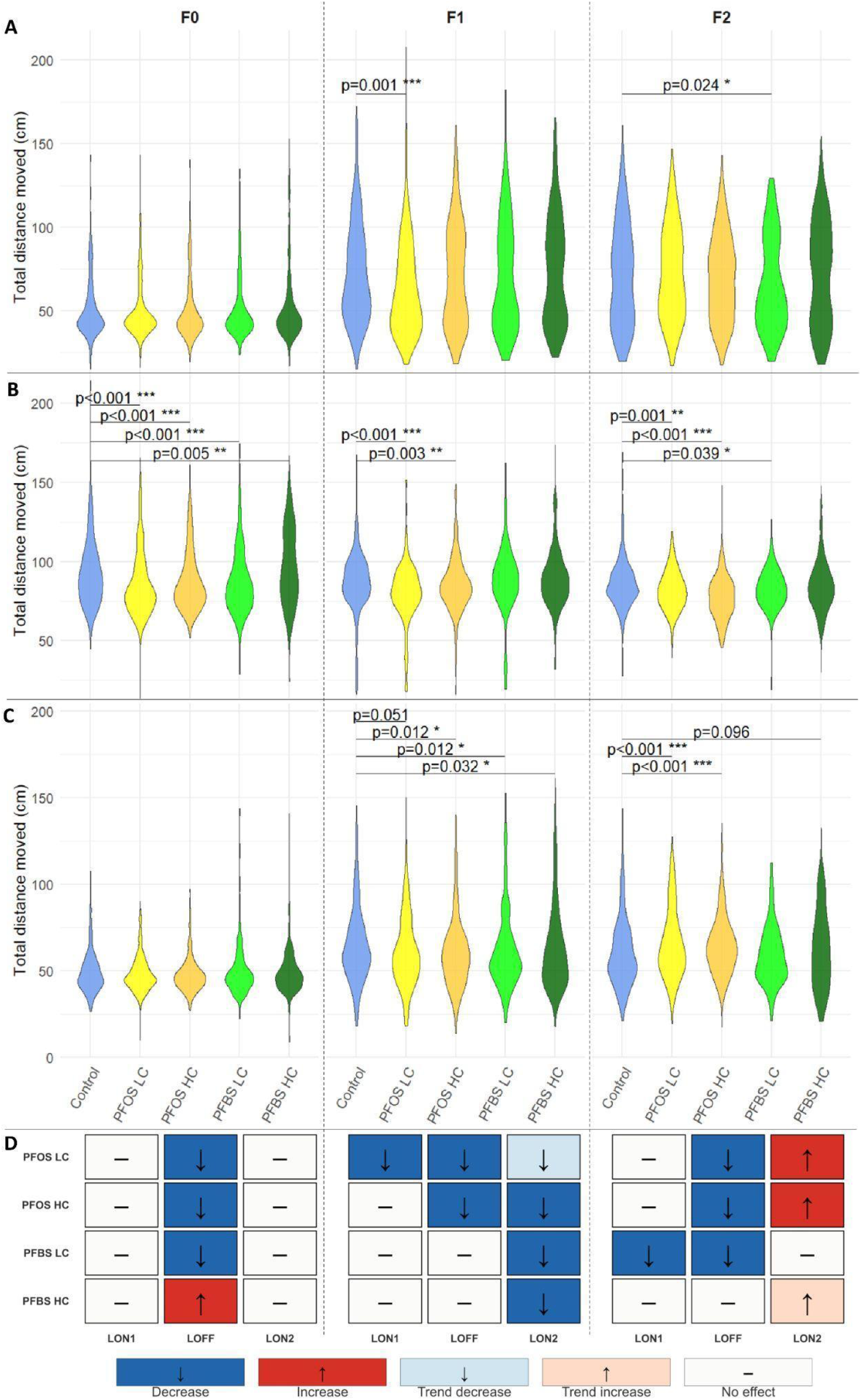
Developmental PFOS or PFBS exposure alters larval photomotor response across three generations. The total distance moved by larvae at five days post-fertilization (5 dpf) is shown for three sequential light phases: (**A**) LON1 (acclimatization). (**B**) LOFF (dark stimulus). (**C**) LON2 (recovery). Within each phase, the five exposure groups are represented as follow: Control (blue), PFOS LC (yellow), PFOS HC (orange), PFBS LC (light green), and PFBS HC (dark green), and are displayed for the F0, F1, and F2 generations (left, center, right columns for each phase respectively). Significant differences from wilcox robust comparisons (WRS2 package) using 20% trimmed means with Dunnett-style multiple comparisons and *p*-values are represented on each panel with significance level. (**D**) Summary of altered behaviors for each generation, relative to their respective control groups. Hypoactivity is indicated in blue, while hyperactivity in red.

In F1, we observed hypoactivity for PFOS LC compared to the control group. This hypoactivity was consistent throughout all phases of the test, including the period before the stressor (LON1 adjusted p-value = 0.001; LOFF adjusted p-value < 0.001; and a trend at LON2 with adjusted p-value = 0.051). F1 PFOS HC also exhibited hypoactivity during the dark phase - LOFF (adjusted p-value = 0.003) and in LON2 (adjusted p-value = 0.012). F1 PFBS LC and HC also showed altered behaviors, with hypoactivity occurring during LON2 (LC adjusted p-value = 0.012 and HC adjusted p-value = 0.032).

In the F2 generation, PFOS LC and HC larvae showed no significant alteration during LON1, however, we observed hypoactivity during LOFF (LC adjusted p-value = 0.001, HC adjusted p-value < 0.001) and hyperactivity in the LON2 phase (LC adjusted p-value < 0.001, HC adjusted p-value <0.001). In comparison, F2 PFBS LC showed hypoactivity during LON1 and LOFF (LON1 adjusted p-value = 0.024; LOFF adjusted p-value = 0.039) while at high concentrations of PFBS, we observed no significant changes in the LPMR test.

These results demonstrate that both treatments in at least one concentration induced transgenerational effects, most prominently seen as hypoactivity in the LOFF phase.

### Developmental PFAS exposure induces multi- and transgenerational transcriptional changes

Following the phenotypic assessment, we examined the multigenerational effects of PFOS and PFBS on gene expression in zebrafish larvae at 5 dpf, the same developmental stage used for behavioral testing. Of note, the larvae used for transcriptomic analysis were not used for behavioral assays, thereby reflecting baseline gene expression alterations in the absence of light-induced stressors.

Overall, our results revealed a greater impact of PFOS and PFBS on the F0, with a maximum of 1637 differentially expressed genes (DEGs) in PFOS HC and a minimum of 795 DEGs in PFBS LC. In F1, only 2 genes were differentially expressed in the PFOS LC and PFBS HC groups, and 3 to 5 genes were identified in each of the other experimental groups. In contrast, F2 exhibited a larger number of DEGs, with a minimum of 7 genes in the F2 PFOS LC group, indicating that this group was generally the least affected in terms of transcriptomic changes, while a maximum of 388 genes was found for the F2 PFBS HC group (Fig. 2A and table S1).

**Fig. 2.**
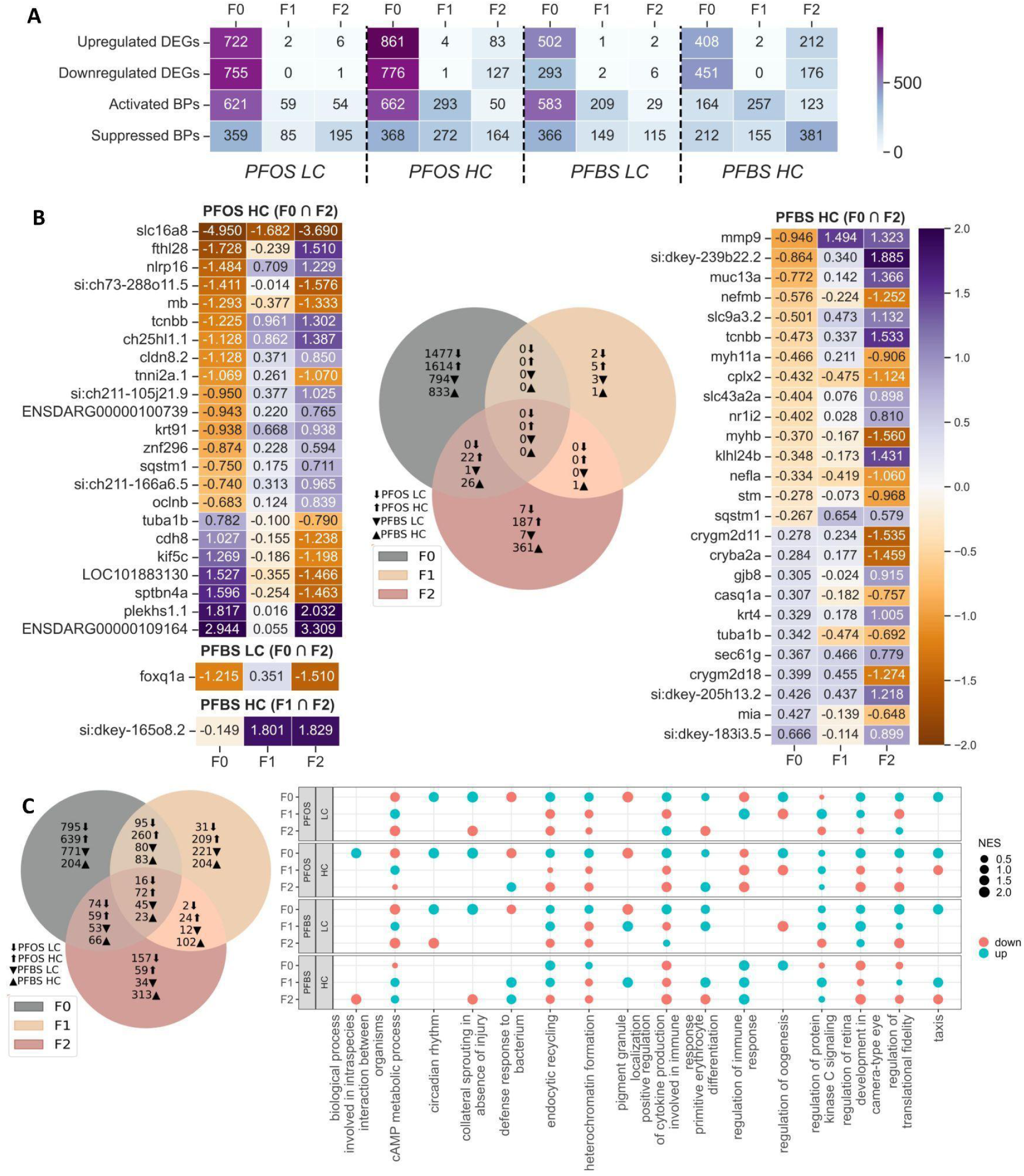
Gene expression changes induced by PFOS and PFBS. (**A**) Heatmap showing the number of significant DEGs and Gene Ontology: Biological Process (GO:BP) terms (*adjusted p-value* < 0.05) across exposure conditions and generations. (**B**) Venn diagram showing the shared and unique DEGs among the three generations, alongside heatmaps presenting log fold-change in expression of shared DEGs between any pair of generations where available. Note that relative expression of all generations is shown although the changes were only significant in F1 and F2 for si:dkey-165o8.2, and in F0 and F2 for the rest. (**C**) Venn diagram illustrates the common and specific GO:BP terms associated with the DEGs across the three generations. (**D**) Dot plot presenting rrvgo-based reduction of GO:BP terms. A semantic similarity threshold of 0.99 was applied to group highly redundant terms and retain representative (Parent) GO:BP terms across different generations and exposure conditions. Each dot represents a gene set. Dot size corresponds to the absolute Normalised Enrichment Score (absNES), indicating the strength of enrichment. Dot color reflects the direction of enrichment. F0, exposed during development - from fertilization to the end of the larval stage (28 dpf); F1, directly exposed as germline; and F2, unexposed.

To evaluate multigenerational persistence of gene expression alterations, we compared DEGs across F0, F1, and F2 (Fig. 2B). We observed no significant DEG overlap across all the three generations in any of the exposure groups; however, F0 and F2 showed overlap in all exposure conditions except PFOS LC. For the HC exposures, 23 and 26 DEGs were shared between F0 and F2 in PFOS and PFBS respectively. Within PFOS’ 23 shared DEGs only four genes were down-and two genes consistently up-regulated between these two generations. For PFBS HC, six were down- and five consistently up-regulated in PFBS (Fig. 2B).

To assess which pathways the differentially expressed genes may be implicated in, Gene Set Enrichment Analysis (GSEA) was employed across the entire gene list using their respective Wald test statistics. GSEA identified a number of affected biological processes (BPs), molecular functions (MFs) and cellular components (CCs) across all conditions (Fig. 2A and table S2). To evaluate the lists of terms and compare these more readily between the exposures, parental terms were identified and organized into clusters, and the most distinct term within each cluster was selected to serve as its representative. Sixteen, 72, 45, and 23 parent (representative) BPs were consistently affected across generations for PFOS LC, PFOS HC, PFBS LC and PFBS HC, respectively (Fig. 2C). BPs across all exposures and generations were related to, among others, regulation of cellular signalling, function, and responses, developmental processes such as oogenesis and eye development, as well as circadian rhythm (Fig. 2C and table S3).

In conclusion, all treatments affected gene expression, predominantly in the F0 generation, with marginal consistency across generations but a discernible overlap between F0 and F2 at HC.

### Developmental PFAS exposure mediates multi- and transgenerational DNA methylation changes

To investigate epigenetic changes potentially underlying the observed multigenerational effects, we analysed differences in 5-methylcytosine (5mC) between F0, F1, and F2 exposure and control groups. DNA extracted from the same 5 dpf larvae as for transcriptome analyses was subjected to Oxford Nanopore sequencing to cover the whole genome and avoid chemical modifications of the DNA prior to the analysis. The samples showed variable depth but sufficient genome-wide representation. Coverage was comparable across F0 and F1 samples, whereas F2 exhibited greater heterogeneity, with several samples showing reduced genomic coverage (fig. S2 and table S4). For the covered CpGs in all conditions, a slight but statistically significant (χ² test, p<0.05) overrepresentation in intronic and intergenic regions, and an underrepresentation in promoters, 5′ untranslated regions (5′UTRs), 3′UTRs, and exonic regions was observed as compared to the CpG relevant genome (fig. S3), which may reflect a potential bias in Oxford Nanopore Technology.

Comparison between exposed and unexposed groups in each generation identified a large number of differentially methylated positions (DMPs). These were subsequently linked to the nearest gene or closest transcriptional start site (TSS), defining differentially methylated genes (DMGs) (Fig. 3A and table S4). Overall, we observed effects across all three generations. Direct exposure to the chemicals (F0 generation) resulted in the highest number of DMPs with PFBS LC showing the strongest effect across 76551 DMPs linked to 17070 genes. Effects of exposure appeared to have propagated to the F1 generation, showing high numbers of DMPs although slightly lower as compared to F0. In F2, the effect could still be observed with thousands of DMPs identified across both concentrations of PFOS and PFBS (Fig. 3A). No trend towards global hypo- or hypermethylation was observed in any of the groups, with the amount of hypermethylated sites in the groups ranging from 43.80% to 54.21% (Fig. 3B and table S5).

**Fig. 3.**
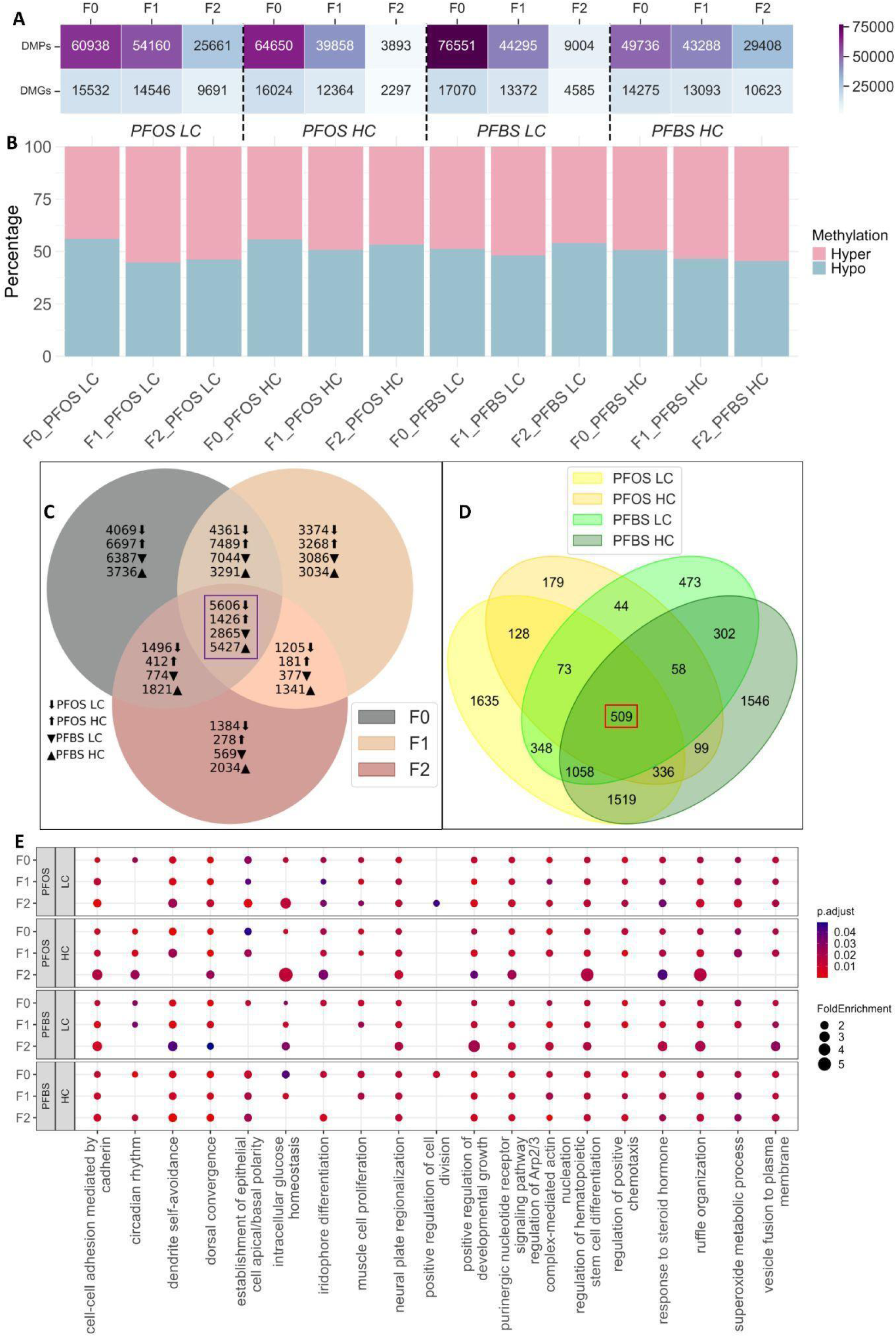
Differential methylation patterns across generations, exposure groups, and concentrations. (**A**) Number of DMPs and DMGs across all generations, exposures, and concentrations. Color intensity reflects the magnitude of changes, with the spectrum ranging from light blue, representing the least affected group, to purple for DMPs and DMGs at the highest values. (**B**) Percentage of hypermethylated and hypomethylated differentially methylated positions (DMPs) for each group. (**C**) Venn diagrams illustrating the number of shared DMGs across different generations for each exposure group. (**D**) Venn diagram showing the overlap of shared DMGs across generations for each exposure condition, corresponding to the regions highlighted in purple square in **C**. In total, 509 DMGs, indicated with a red square, were commonly identified across all exposure conditions and generations. (**E**) Dot plot showing rrvgo-based reduction of GO:BP terms of DMGs. A semantic similarity threshold of 0.95 was applied to group highly redundant terms and retain representative (Parent) GO:BP terms across different generations and exposure conditions. Fold enrichment quantifies the extent to which terms are observed more (or less) frequently than expected by chance. A fold enrichment greater than 1 indicates that the terms are overrepresented. The enriched terms are significant (adjusted p-value < 0.05), as indicated by the red-colored dots.

To investigate whether DMPs persisted across generations, we analysed the overlaps between F0, F1, and F2 within each exposure group. While overlap of DMPs across generations was limited (fig. S4), gene-level aggregation revealed shared DMGs, suggesting convergence at the gene level, despite site-specific variability. Indeed, thousands of DMGs overlapped, with the highest number (5606) in PFOS LC and the lowest (1426) in PFOS HC (Fig. 3C). Notably, 509 DMGs were consistently shared across all exposure conditions and generations (Fig. 3D). A detailed list of these shared DMGs is provided in table S6.

Subsequently, we conducted enrichment analysis of all DMGs (table S7) and assessed persistence of altered BPs resulting in 212, 48, 89, and 198 BPs overlapping across generations for PFOS LC, PFOS HC, PFBS LC, and PFBS HC, respectively (fig. S5). BPs linked to DMGs across all generations and exposure groups included terms related to circadian rhythm, brain function, such as dendrite self-avoidance and signaling pathway, and regulatory processes, including hematopoietic stem cell differentiation, chemotaxis, cell division, actin nucleation and developmental growth (Fig. 3E and table S8).

### Correlation between expression and DNA methylation implies functionality of PFAS induced changes

Given the relationship between methylation and gene expression, we assessed the overlap between DEGs and DMGs, hereafter referred to as DEG-DMG pairs. Our analysis revealed overlaps in the F0 ranging from 374 genes in PFBS HC to 1034 genes in PFOS HC. In comparison, these overlaps were much lower in the F1 and F2 generations, reflecting the lower numbers of DEGs in these generations (Fig. 4A). No DEG–DMG pairs were consistently overlapping across all three generations in any exposure condition. However, we identified a transgenerational overlap between F0 and F2, with one (*sptbn4a*) and four (*nr1i2, myh11a, cplx1, klhl24b*) shared DEG–DMG pairs in the PFOS HC and PFBS HC groups, respectively. Across all DEG-DMG pairs, we observed a broadly similar distribution over the four possible methylation–expression relationships (hypermethylation + down-regulation, hypomethylation + up-regulation, hypermethylation + up-regulation, hypomethylation + down-regulation) (fig. S6).

**Fig. 4.**
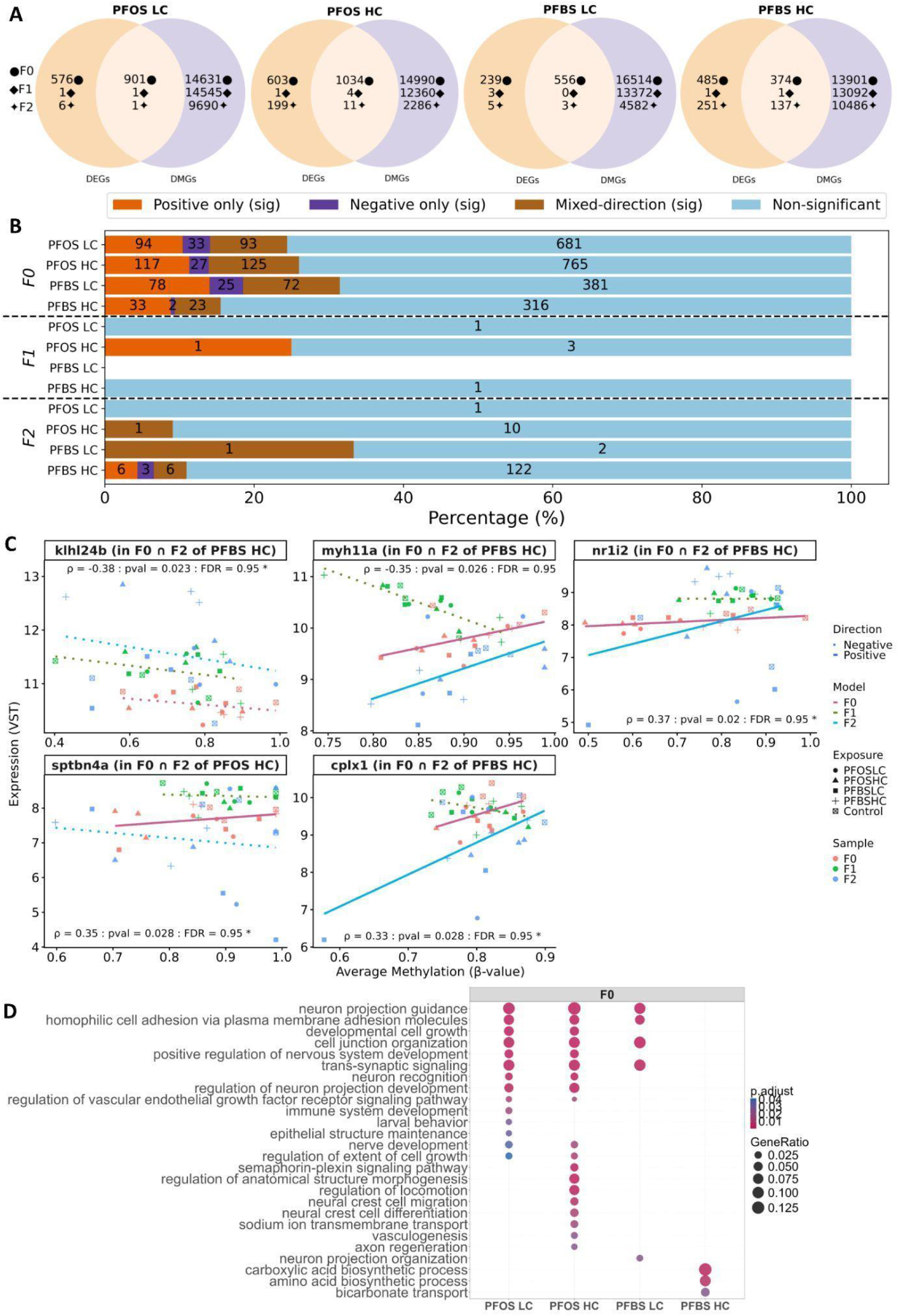
DEG-DMG pairs reveals methylation–expression coupling. **(A**) Overlap between DMGs and DEGs across exposure groups and generations. **(B)** Number and directionality of DEGs-DMGs correlations: significant (p<0.05) positive correlation “positive (sig.)”, significant negative correlation “negative (sig.)”, genes with DMPs showing significant positive and negative correlation “mixed-direction (sig)**”** and Non-significant correlation. **(C)** Pairwise scatter plots showing correlations among five DEG-DMG pairs overlapping between F0 and F2, in the PFOS HC and PFBS HC. Each panel represents the relationship between the average β-value across all DMPs of each gene and their corresponding gene expression levels, with points indicating individual samples. The median Spearman correlation coefficient (ρ), median p-value, and median FDR across all DMPs associated with each gene were also presented. Dotted lines denote the linear regression fit across samples. All five genes contain loci with both positive and negative correlations indicated as mixed-direction (*). **(D)** Dot plot showing rrvgo-based reduction of GO:BP terms significantly correlated DEG-DMG pairs using a semantic similarity threshold of 0.7 to group redundant BP terms and retain representative BPs across F0 exposure conditions. Gene Ratio indicates the proportion of DEG-DMG pairs associated with the representative BP, with higher values indicating stronger representation in the gene set. Dot color represents the significance of enrichment (adjusted p-value < 0.05).

To assess whether DEG-DMG pairs reflect functional regulatory relationships, we performed correlation analyses. To this end, we integrated data from all exposure groups across generations and conducted pairwise Spearman rank correlations between locus-level CpG methylation and their associated gene expression values in matched samples. The resulting correlation profiles revealed multiple CpG–gene pairs exhibiting statistically significant (FDR < 0.05) correlations (table S9, A and B). These findings indicate that a subset of exposure-associated methylation changes is correlated with concordant alterations in gene expression, supporting their potential regulatory relevance. Next, we overlapped the correlation results from the integrated data with the DEG-DMG pairs. As multiple testing correction resulted in few DEG-DMG pairs at FDR < 0.05, we used a nominal p-value threshold (p < 0.05) to retain additional DEG-DMG pairs for exploratory downstream analyses. This comparison identified a subset of DEG-DMG pairs exhibiting significant (p<0.05) methylation–expression correlations (table S10). Both negative and positive correlations were observed, consistent with context-dependent regulatory effects of DNA methylation on gene expression. While some DEG-DMG pairs showed exclusively significant positive or negative correlations, others contained both positively and negatively correlated DMPs (Fig. 4B). For one (*sptbn4a*) and four (*nr1i2, myh11a, cplx1, klhl24b*) DEG-DMG pairs exhibiting transgenerational overlap between F0 and F2 in PFOS HC and PFBS HC, respectively, expression and methylation was significantly correlated (Fig. 4C). Yet, the proportion of significantly correlated DEG-DMG pairs did not exceed 40% across all exposure groups, likely reflecting the use of whole-larvae samples in this analysis.

To further assess the functional relevance of the significantly correlated (p<0.05) DEG-DMP pairs, we performed GO enrichment analysis for the F0 exposure groups only; F1 and F2 exposure groups were excluded due to insufficient gene counts for reliable statistical testing. For all F0 exposure groups, 139 significantly (FDR < 0.05) enriched BPs were identified (table S11), indicating strong functional organization within these gene sets. To reduce redundancy among enriched GO terms and facilitate biological interpretation, related biological processes were grouped into clusters and 26 representative terms summarizing the dominant functional BPs across the clusters were identified (Fig. 4D). The summarized enrichment results revealed both shared and exposure-specific BPs. Notably, neuron projection guidance, trans-synaptic signaling, homophilic cell adhesion via plasma membrane adhesion molecules and cell junction organization were consistently represented across three (PFOS LC, PFOS HC and PFBS LC) of the four exposure groups, suggesting a common functional component associated with regulation of neuron–neuron adhesion and the assembly of specialized cell junctions required for synapse formation, stability, and efficient synaptic signaling. In addition, regulation of neuron projection development, regulation of growth factor receptor signaling pathway, nerve development, and regulation of the extent of cell growth were enriched in PFOS LC and PFOS HC, suggesting further shared functional roles among these groups in signal transduction regulation and neurodevelopment.

### DNA Methylation changes associate with F2 Behavior

Given the large number of consistent DNA methylation changes across generations, we explored if such changes were associated with F2 behaviors, and thus could potentially underlie transgenerational effects and serve as predictors of such effects. We conducted an association analysis using three models: Model-F0, Model-F1 and Model-F2 predicting F2 behavioral phenotypes using aggregated DNA methylation profiles from all exposure conditions in F0, F1 and F2, respectively. This analysis identified significant (p-value < 0.05) behavior-associated positions (BAPs) (table S8). We reasoned that the positions overlapping between the models could be carriers of transgenerational information and most robust predictors of F2 outcomes. A total of 68, 47, and 69 BAPs shared across the three models were identified in LON1, LOFF and LON2, respectively (Fig. 5, A to C and table S12).

**Fig. 5.**
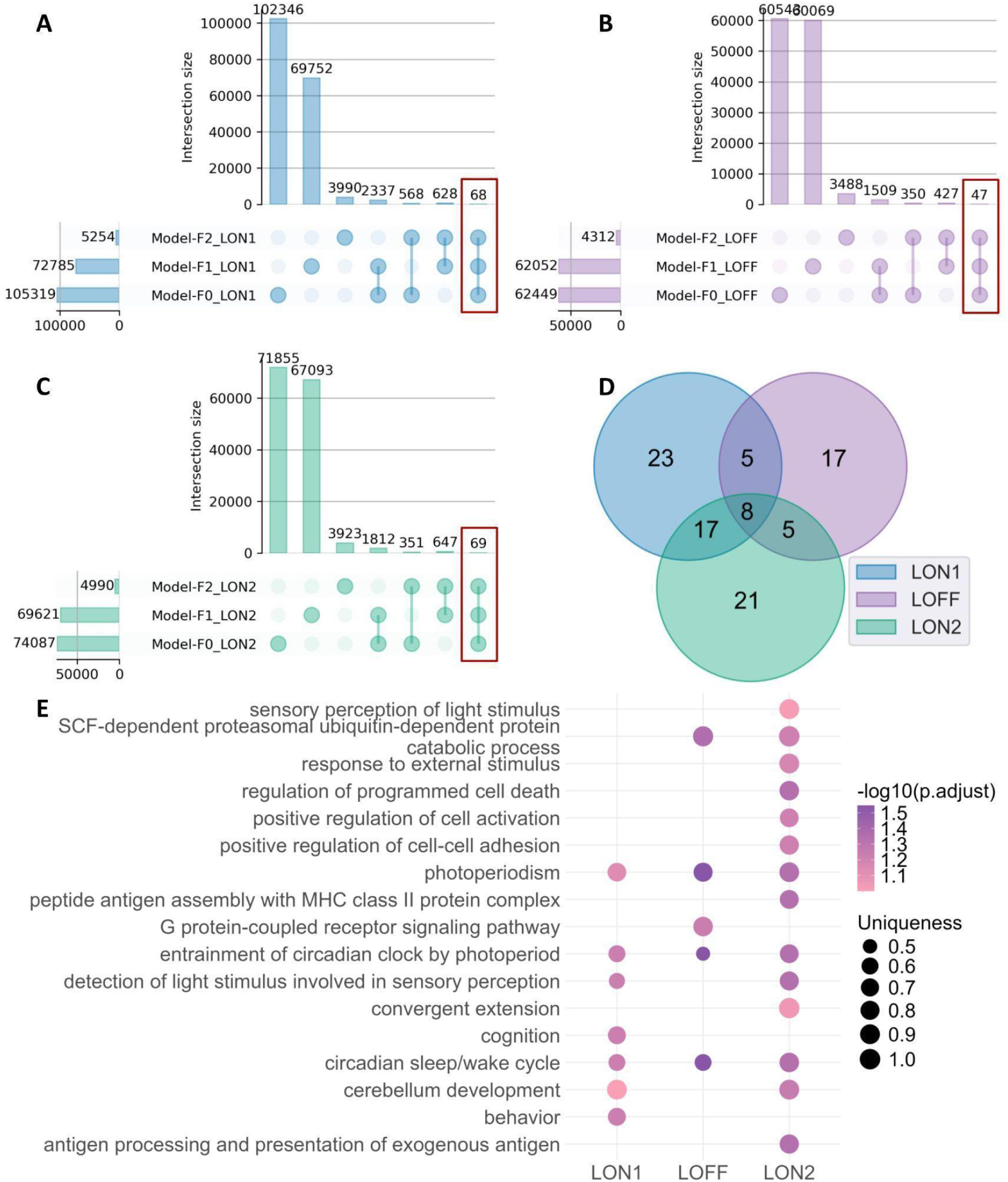
DMPs associated with behavior in F2. (**A to C**) Number of BAPs and overlaps (highlighted in red) between models in LON1, LOFF and LON2 conditions. Model-F0, Model-F1 and Model-F2 predict F2 behavioral phenotypes using aggregated DNA methylation profiles from all exposure conditions in F0, F1 and F2, respectively. (**D**) Overlap between genes associated with behavior-associated positions (BAP-genes) across the different light phases of the LMPR test. (**E**) Dot plot summarizing enriched GO:BP parent terms at LON1, LOFF, and LON2; parent terms were identified using rrvgo clustering of enriched BPs with a similarity threshold ≥ 0.5 (range: 0–1; lower values yield broader clusters). Only terms with a dispensability score of 0 (range: 0–1; higher values indicate more dispensable, i.e., redundant, terms) were retained as parent terms (table S12). Dot size represents the GO:BP uniqueness, and color indicates the –log₁₀ of the Benjamini-Hochberg false discovery rate (FDR).

To determine the functional significance of the identified BAPs, these sites were first mapped to their nearest genes. This resulted in 53, 35, and 51 genes linked to behavior-associated positions (BAP-genes) for LON1, LOFF, and LON2, respectively (table S12). Eight BAP-genes (*Fbxl3a*, *si:ch211-207l22.2*, *si:ch73-116o1.3*, *si:dkey-183k8.1*, *si:ch211-287c11.2*, *si:dkeyp-4c4.2*, *ENSDARG00000098628* and *ENSDARG00000114602*) overlapped between light conditions (Fig. 5D). For the only coding gene of these eight, *Fbxl3a*, expression and DNA methylation was inversely correlated in all generations, suggesting a functional role of DNA methylation levels at the identified position (fig. S7). Fbxl3a exhibits a conserved function in circadian rhythm generation. Subsequent enrichment analysis of all BAP-genes also revealed significant (FDR < 0.1) associations with functions related to the circadian rhythm as well as pathways related to brain and nervous system functions (Fig. 5E).

## DISCUSSION

Despite concerns over PFAS chemicals in the environment, the extent and mechanisms of their multigenerational effects remain poorly characterized. This study investigated the multi- and transgenerational impact of PFOS and its replacement, PFBS. We assessed whether these chemicals cause behavioral effects that persist beyond direct exposure and aimed to identify the potential molecular mechanisms involved, as well as whether molecular changes induced by direct exposure can be used to predict effects across generations. To this end, we exposed the F0 generation at environmentally relevant concentrations and examined the subsequent offspring. Our results demonstrate that exposure to PFOS and PFBS induces multigenerational behavioral effects and that these changes co-occurred with alterations in gene expression and DNA methylation associated with processes relevant for development and functioning of the nervous system. In addition, some of the epigenetic changes observed in the F0 generation were associated with behavioral outcomes in the F2 generation.

The LPMR assay showed that developmental PFOS and PFBS exposure at environmentally relevant concentrations altered locomotor activity across F0-F2 with both hypo- and hyperactivity depending on generation, concentration, and light phase. These effects were most evident in the LOFF (stress) and LON2 (recovery) phase than in the LON1 (basal condition), suggesting that early-life PFAS exposure specifically impairs behavioral response to stress rather than baseline activity.

In the F0 generation, PFBS effects were less consistent across generations, whereas PFOS exposures caused hypoactivity in the LOFF phase, persisting to both F1 and F2. Previous behavioral studies using zebrafish have also identified the LOFF phase as particularly sensitive to neuroactive pollutants, probably due to its dependence on the rapid integration of photic stressors (*32*). In contrast, a study by Haimbaugh et al. (*23*) reported hyperactivity, which was only observed in F1 and F2 generations, following a shorter exposure to PFOS, limited to the first 5 dpf. Of the three different concentrations (7, 70 and 700 ng/L), transgenerational effects were observed only at 70 ng/L, suggesting a narrow and concentration-specific window for heritable behavioral disruption in their study. Such discrepancies may arise from differences in exposure concentration and length. Our 28-days exposure spanned over the embryonic and late larval stages, including periods of neuronal circuit maturation (*33*), glutamatergic and dopaminergic neurons maturation (*34*–*36*) and optic tectum development (*37*). Additionally, it coincided with later stages of sex differentiation and germ cell development which include epigenetic reprogramming and might have reinforced the heritability of the phenotype (*38*).

For PFBS, LOFF-responses were inconsistent across generations. In the LON2, we noted a similar pattern for both PFOS exposures and PFBS HC, with no effect in F0, hyperactivity in F1 and hypoactivity in F2. This phenomenon has been observed in other transgenerational studies. For example, in prior work using mice, DTT-induced obesity did not manifest in the exposed parental generation but was observed in the F3 generation (*39*). Biologically, this can be explained by the fact that in the case of direct exposure, chemicals act on somatic cells. In contrast, intergenerational effects occur when chemical exposure, along with outcomes manifesting in F0, impact the germline, which can result in specific phenotypic traits in the F1. For transgenerational effects, the chemical does not directly interact with the affected generation; instead, the impact of the exposure is inherited solely through biological factors (*2*). The reason for the differences between PFOS and PFBS effects in our study might be because PFBS is less bioaccumulative than PFOS (*13*) and may act through partially distinct molecular targets. Nevertheless, the presence of F2 hypoactivity in some PFBS groups suggests that even lower-bioaccumulative PFAS can elicit heritable behavioral phenotypes under certain exposure scenarios.

The observed phase and concentration-specific locomotor effects correspond to the axonogenesis, neuronal signaling, and visual system development pathway in both DEG and DMG analyses.

In terms of transcriptomic changes, the most pronounced alterations were observed in the F0 generation, followed by F2 and then F1. In fact, in F1, we identified only a few DEGs, this might reflect actual biological responses. While DEGs varied across generations, subsets were shared between F0 and F2, particularly in the high concentration groups. Lack of overlap with F1 DEGs is most likely due to the small number identified in this generation. In the PFOS HC, 23 DEGs were common to F0 and F2. These included genes involved, or predicted to play a role, in cytoskeletal organization and neuronal functions, such as *tuba1b* and *sptbn4a* (*40*, *41*), or *sqstm1* and *kif5c*, which are associated with key neuronal processes, including axonogenesis and synaptic vesicle transport, respectively (*42*, *43*). Mutations in *kif5c* have been associated with brain malformations and infantile onset epilepsy in humans (*44*), while studies in mice support an important role for *Kif5c* in the cortical development and maintenance of motor neurons (*45*, *46*), which may contribute to behavioral alterations. In the PFBS HC group, 26 DEGs were shared between F0 and F2. These included *tuba1b* and *sqstm1*, as well as neurofilament light chain a (*nefla*) and neurofilament medium chain b (*nefmb*), both involved in the neurofilaments, essential components of axons. Notably, this group also included several crystallin genes (*cryba2a, crygm2d11, crygm2d18*), which are key structural components of the eye lens. This might suggest alterations in visual systems that could influence behavior in the light/dark test response.

Although the DEGs differed between conditions, many were associated with overlapping biological pathways among F0, F1, and F2. For PFOS (both LC and HC) and PFBS HC, enriched pathways were consistently related to visual perception and brain development and function, including axonogenesis and synaptic signaling. These results are in line with a previous study showing that direct exposure to PFOS affects eye morphology (*47*). However, PFBS LC presents a different case, since despite inducing significant hypoactivity in F0, F1, and F2, it did not show enrichment of pathways related to the brain or vision. Thus, the behavioral changes observed upon PFBS LC might be due to other mechanisms that we could not pinpoint via transcriptional analyses.

Epigenetic changes are associated with the development and inheritance of specific phenotypes, and prior studies have identified epigenetic and transcriptional alterations in epigenetic modifiers upon PFAS exposure, e.g., in histone lysine-specific demethylases in mice directly exposed to PFOS (*48*) and in *dnmt* genes in zebrafish exposed to PFOS and PFBS (*31*). Therefore, in addition to gene expression analyses, we examined DNA methylation in F0, F1, and F2 fish. Our data revealed extensive DNA methylation changes in groups exposed to PFOS and PFBS in all three generations. Generally, the number of identified DMPs was much higher than in previous transgenerational studies in zebrafish (*31*, *49*)

We assume that this is not due to differences in the used chemicals and exposure schemes but rather in the method for assessing DNA methylation. Previous studies have mainly employed reduced representation bisulfite sequencing (RRBS), while in this study, we made use of the 3rd generation sequencing method ONT. ONT excels by not requiring chemical modification of the DNA, such as bisulfite conversion, and very long reads, thus giving a more appropriate representation of DNA methylation (5-methylcytosine) differences induced by the exposure.

The identified DMPs were found in or near genes associated with neurodevelopment, including in the PFBS LC group, where this pattern suggests that, despite the lack of corresponding transcriptional changes, neurodevelopmental processes may still be affected. Among differentially methylated genes in both PFOS and PFBS, we identified *irx1a, nrxn2a,* and *cerkl*. The methylation of *irx1a* human ortholog, *IRX1*, has been shown to correlate with the efficiency of neuronal differentiation in human iPSCs (*50*). Alterations in *nrxn2a* expression have been connected to differences in locomotor activity in zebrafish larvae (*51*), while *cerkl* is recognized for its role in visual disorders and is crucial for retina development in zebrafish embryos (*52*). Shared BPs across generations, and both chemical treatments, included axonogenesis, axon development and guidance, synapse formation, and signaling pathways involved in the formation of axons and neural circuits (tyrosine kinase receptor signaling and cell-cell adhesion).

Additionally, we found considerable overlap between differentially methylated and differentially expressed genes in F0. In fact, around half of the DEGs identified in F0 overlapped with DMGs for all exposure conditions. On the other hand, hardly any overlaps were detected in F1 and F2. This could imply that the methylation changes in F0 are directly translating into expression changes, alternatively that transcriptional alterations result in epigenetic changes in F0. In contrast, in the other generations, the detected DNA methylation changes are not affecting transcription of the nearest genes at 5 dpf. Notably, there were about equal numbers of directly (hypomethylation - downregulation / hypermethylation -upregulation) and inversely (hypomethylation - upregulation / hypermethylation - downregulation) related overlapping genes. This can be explained by the fact that most of the DMPs did not lie in the promoter region, and it is known that for other genomic contexts, DNA methylation does not always play a repressive role (*53*).

Correlation analysis integrating DNA methylation and gene expression revealed a subset of DEGs associated with DMGs, suggesting potential epigenetic regulation of transcriptional responses across exposures. Indeed, a number of DEG-DMG pairs showed significantly correlated levels of DNA methylation and expression. Unlike F1 and F2, which generally resulted in none or very few shared DEGs–DMGs, F0 yielded more significantly correlated DEG–DMG pairs across all exposure groups, which are enriched in pathways related to neurodevelopment, indicating that exposure-associated methylation changes may contribute to transcriptional modulation of genes involved in neural development and function. These findings support the hypothesis that environmentally induced DNA methylation alterations can be linked to coordinated gene expression changes, consistent with previous studies showing that environmental exposures can modify epigenetic marks that regulate transcriptional programs *(77-80)*. Ubiquitin-conjugating enzyme E2Ka *(ube2ka*), the only significantly correlated DEG-DMG pair identified in F1, has been implicated in the regulation of chromatin dynamics via ubiquitination and proteasome-mediated turnover of histone H3. Through this mechanism, it can influence transcriptional repression marks and downstream gene expression programs, suggesting that the observed methylation–expression correlation may reflect exposure-related modulation of epigenetic regulatory processes *(81)*.

Among the significantly correlated DEG-DMG pairs identified in F2, transcription factor AP-2 gamma (*tfap2c*) stands out as a transcription factor with well-documented roles in regulating multiple developmental and signaling genes *(82-86).* Epigenetic modification of *tfap2c* may influence key transcriptional networks, particularly in developmental contexts. Similarly, *nr1i2* is particularly notable as it encodes the pregnane X receptor (PXR), a nuclear receptor transcription factor that regulates xenobiotic metabolism and detoxification pathways *(87,88).* Exposure-associated methylation changes affecting *nr1i2* may influence transcriptional responses to environmental chemicals *(89)* such as PFAS, consistent with evidence that epigenetic modification of *nr1i2* regulates such transcriptional activity *(90,91)*. Genes such as *chl1a (92), slit3 (93,94), cdh26.1 (95), pcdh9 (96-99)*, and *prnpb (100,101)* participate in signaling pathways and structural networks that feed into transcriptional regulation, especially in neural and developmental pathways. Several recent studies have associated epigenetic dysregulation of these genes with possible implications for disease pathogenesis *(102)*. Although no DEG-DMG pairs were shared across all generations, a subset overlapped between groups F0 and F2, namely *sptbn4a, nr1i2, myh11a, cplx1*, and *klhl24b*. Notably, *nr1i2* (discussed above) was among the overlapping genes, while the remaining ones participate in neuronal signaling, stress responses, cytoskeletal organization, and ubiquitin-mediated protein regulation, biological processes known to be sensitive to epigenetic modification *(103-105)*.

Overall, the integration of DNA methylation and transcriptional data provides evidence that a subset of exposure-associated epigenetic modifications may be functionally linked to gene expression regulation, particularly within neurodevelopmental pathways in F0. However, although we found significant correlation for some of the DEG-DMP pairs, we cannot exclude the possibility that these are coincidental and lack functional significance at this developmental stage. Another important consideration is that our analysis was conducted on whole larvae. As such, methylation changes that occur in a specific subset of cells may not be accompanied by detectable transcriptional changes at the whole-organism level, potentially masking cell-type specific regulatory effects. Thus, it remains to be determined whether the identified methylation changes actively regulate gene expression and contribute directly to the observed phenotypic outcomes.

While epigenetic changes may differ between the directly exposed F0 and unexposed F2 generations due to the different ways chemicals can alter phenotype, the molecular alterations in F0 may initiate biological cascades that influence or reflect mechanisms leading to phenotypic changes in descendants. Given the evidence that the majority of exposure groups exhibited both behavioral and DNA methylation transgenerational alterations, we investigated whether epigenetic modifications in the F0 generation could serve as predictors of behavioral outcomes in the unexposed F2 generation. Using integrated epigenetic and behavioral datasets, we developed a model to identify positions that are differentially methylated in all generations and associated with locomotor activity in F2. We found 68, 47, and 69 DMPs associated with behavior during LON1, LOFF, and LON2 phases of the test in F2, respectively. Eight genes were consistently identified for LON1, LOFF, and LON2 (*fbxl3a*, *si:ch211-207l22.2*, *si:ch73-116o1.3*, *si:dkey-183k8.1*, *si:ch211-287c11.2*, *si:dkeyp-4c4.2*, *ENSDARG00000098628*, *ENSDARG00000114602*). Notably,

*fbxl3a*, the only coding gene of eight in common for all three conditions, is involved in circadian regulation in zebrafish (*54*). A link to circadian function was also found in the pathways linked to all DMGs associated with F2 behaviour, including circadian sleep/wake cycle, entrainment of circadian clock by photoperiod, detection of light stimulus involved in sensory perception, photoperiodism, sensory perception of light stimulus, response to external stimulus, and G protein-coupled receptor signaling pathway. A number of chemicals, including PFAS (*55*) have been shown to affect circadian rhythm. Furthermore, PFOS, but not PFBS, was shown to affect the visual system in zebrafish (*56*). However, these effects so far have not been investigated in transgenerational settings. It remains to be clarified if changes in light perception and/or circadian rhythmicity underlie the behavioral effects described in this study.

While this study provides novel insights into the multigenerational effects of PFOS and PFBS and underlying epigenetic changes, several limitations must be acknowledged. Firstly, this study has not established causality between phenotypic and molecular changes. Future studies employing epigenome editing will shed more light on causal involvement of the identified DNA methylation changes. Moreover, the DNA methylation changes in the F0 generation associated with behavioral phenotypes in the unexposed F2 generation require validation using independent datasets generated with comparable methodologies. Unfortunately no such dataset could be identified and will need to be developed in the future. Another limitation is the focus on DNA methylation while other epigenetic modifications or mechanisms are likely involved in the multi-and transgenerational phenotypes.

In summary, this study demonstrates that developmental exposure to both PFOS and PFBS induces persistent behavioral changes in zebrafish extending beyond the directly exposed generation. These phenotypes were accompanied with transcriptomic and DNA methylation alterations, some of them persistent across all generations. This study also identified DNA methylation changes induced in the F0 generation that were associated with behavioral outcomes in the unexposed F2 generation. This suggests that DNA methylation patterns could serve as early biomarkers for predicting transgenerational effects of PFAS exposure and that our bioinformatic approach is a promising tool to anticipate long-term impacts of environmental contaminants across generations.

## MATERIALS AND METHODS

### Selection of PFOS and PFBS Exposure Concentrations

In the present study, we used PFOS (Potassium salt C_8_HF_17_O_3_SK, CAS 2795-39-3, molecular weight 500.13 g/mol, purity > 98%) and PFBS (Tetrabutylammonium salt (C_4_H_9_)_2_N(C_4_HF_9_O_3_S), CAS 108427-52-7, molecular weight 300.10 g/mol, purity > 98%). Both chemicals were purchased from Sigma-Aldrich (St. Louis, Missouri, USA). We prepared Stock solutions in aquarium water (see 2.2 for more details) and stored at room temperature (1050 µM for PFOS and 825 µM for PFBS). Working solutions were prepared from stock solutions and their concentrations were verified using liquid chromatography-mass spectrometry (LC-MS) and were determined at 2.41 µM for PFOS and 2.64 µM for PFBS.

PFAS concentrations selected to avoid fish lethality and allow a chronic exposure of 28 days. The no-effect-concentration (NOEC) for PFOS was determined using a simplified semi-static Fish Embryo acute Toxicity testing (FET) in accordance with OECD Test Guideline 236 as well as supporting literature. Briefly, we exposed zebrafish embryos from1-hour post fertilization (hpf) for five days in 96 well plates to a range of PFOS concentrations (0.75 nM to 6.00 nM). Lethality was assessed at 144 hpf based on coagulation, lack of somite formation, non-detachment of the tail, and lack of heartbeat. These lethal endpoints were monitored daily from 24 to 144 hpf and compared to unexposed embryos. The result showed an upper NOEC value of 3.716 nM (2 µg/L). This value was designated as the High Concentration (HC); whereas the Low Concentration (LC) was 10 times lower, resulting in a LC of 0.37 nM (0.2 µg/L). For ease of the experiment, similar concentrations were used for PFBS (0.67 nM (0.2 µg/L) for LC and 6.6 nM for HC (2 µg/L)).

### Zebrafish Husbandry and Exposures

Adult zebrafish (*Danio rerio*, AB strain; ZFIN ID: ZDB-GENO-960809-7) were purchased from Karolinska Institute (Stockholm, Sweden) and maintained at the research facilities of Man-Technology-Environment research center (MTM), Örebro University. Husbandry and reproduction followed standard protocols as outlined by Westerfield (*57*). Stockfish used to produce the exposed generation were kept in a recirculating system at 26±1°C with a 14:10h light:dark cycle. Aquarium water was made of tap water filtered through a reverse osmosis device. Water parameters, including conductivity (350 µS), pH (7.2), and temperature (26±1°C), were maintained using a Profilux 3.1 controller (GHL, Advanced Technology). Carbon hardness, nitrates, nitrites (Easy Test strips, JBL), and ammonia (Test NH3/NH4+, Tetra) were monitored weekly. Fish were fed twice daily with TetraRubin® flakes (Tetra, Germany) in the morning and with freshly hatched artemia nauplii (Ocean Nutrition®) in the afternoon. To produce F0 zebrafish embryos, six breeding groups (3 males and 2 females each) were established the day before spawning in 1.4 L spawning tanks (Techniplast, Italy) using stockfish. The next morning, embryos were collected approximately one hour post-fertilization. Seventy fertilised embryos were promptly transferred into 2 L pre-warmed tanks containing the respective exposure solutions. Thus, embryos were chronically exposed to different concentrations of PFOS or PFBS (low concentration (LC) = 0.2 µg/L or high concentration (HC) = 2 µg/L) starting at 1 hpf (hours post-fertilization) and for 28 days. Exposure tanks were kept in an incubator at 26 ± 1 °C, on a 14:10h light:dark cycle, and constant aeration. To maintain stable concentrations over the course of the exposure, the tanks were kept under semi-static conditions with 20% of each exposure solution being replaced daily. Between 28 and 35 days post fertilization (dpf), exposure solutions were progressively replaced by aquarium water. At 35 dpf, fish were transferred to 4 L polycarbonate tanks (Zebcare, Nederweert, The Netherlands) in the flow-through system at the same conditions described for the stockfish.

The F1 and F2 generations were obtained from F0 fish following the same breeding protocol, but embryos were not exposed to any chemicals and were raised in regular aquarium water. F1 fish were maintained until adulthood (>3 months) and used to generate the F2. Unexposed F0 fish were also used to produce F1 and F2 as control for each generation.

All zebrafish experiments were conducted at the MTM Research Center according to ethical permission (Dnr. 13595-2023).

### Larval Photo Motor Response Test

Behavioral assessments were performed at the MTM, Örebro, Sweden. The Larval Photo Motor Response (LPMR) test was used to evaluate locomotion of zebrafish larvae at 5 dpf under both light and dark conditions and was conducted for F0, F1, and F2 zebrafish larvae. Freshly fertilised eggs (1 hpf) from each parental or grand-parental exposure group were pooled and one embryo fish per well was distributed into 96-well plates (Techno Plastic Products, Switzerland) in 200 µL of either exposure solutions (F0) or aquarium water (F1 and F2). Semi-static conditions were maintained by replacing half of the solution daily. Plates were incubated at 26 ± 1°C under a 14:10h light:dark cycle. At 5 dpf, the LPMR test was conducted using the DanioVision and EthoVision XT10 video tracking system (Noldus Information Technology, Netherlands). The LPMR test consisted of the following 4 steps: (i) a 5-minute acclimatization period of larvae to the device under 10 lx light (not recorded), (ii) a 5-minute control phase at 70 lx light (LON1), (iii) 5 minutes dark phase (< 1 lx; LOFF) to induce a stress response, and (iv) a 5 minutes recovery phase under 70 lx light (LON2). All tests were performed in the afternoon, corresponding to the larvae’s most stable activity period. Four plate replicates were performed across 6 different days, resulting in a total of 384 larvae per condition. For each larva, the total distance traveled (cm) was recorded. Data from dead or unhealthy embryos were removed. The remaining data were analysed using R (v4.3.1). Due to non-normal distributions, heteroscedasticity, and the presence of outliers in the behavioral data, differences in larval response among conditions were assessed using robust statistical methods. Wilcox robust statistics analysis was performed (WRS2 package) with 20% trimmed means and Dunnett-style multiple comparisons was employed to compare each treatment group against the control (p<0.05 for significance and p<0.1 for trends).

### DNA and RNA Extraction

In parallel with the maintenance of adult fish and phenotypic evaluations, 30 larvae per exposure group were collected at 5 dpf for each generation (F0, F1, and F2) in triplicate for molecular analyses. Larvae were humanely euthanized using a benzocaine solution (500 mg/L; Ethyl 4-aminobenzoate, 98%, C₉H₁₁NO₂, CAS: 94-09-7; ThermoFisher, Germany) and transferred to Eppendorf tubes. Residual water was removed and replaced with RNAlater solution, and samples were stored at –80 °C until further processing. For DNA and RNA extraction, samples were resuspended in freshly prepared solution of RLT buffer (Qiagen) and 1% beta-mercaptoethanol, and homogenised using a Bullet Blender® (Next Advance). DNA and total RNA were extracted using the AllPrep DNA/RNA minikit (Qiagen) following manufacturer’s instructions. RNA concentrations were quantified using a NanoDrop 2000c (ThermoScientific). DNA concentration and integrity was evaluated using Qubit (ThermoScientific) and Tapestation (Agilent). DNA Integrity Number (DIN) ranged from 6.7 to 9.8, with an average of 8.4.

### RNA sequencing

RNA-Seq library construction and sequencing were conducted by BGI Genomics (BGI Tech Solutions, Hong Kong). DNBSEQ eukaryotic strand-specific mRNA libraries were constructed and sequenced on a DNBSEQ (MGI Tech) platform, generating 100 base pair paired-end reads. Approximately 24 million paired-end reads were generated per sample on average (fig. S8). Raw reads underwent filtering and quality control processes to remove contaminants, low-quality reads, adapter sequences, and artifacts using SOAPnuke software (*58*) with the following filter parameters: "--minReadLen 100 -l 20 -q 0.4 -n 0.001 --polyX 50 --adaMR 0.25". Only bases with a Phred Score ≥ 20 were selected, indicating a minimum accuracy score of 99%. Higher scores indicate higher confidence in the accuracy of the base call, with a score of 40 indicating 99.99% accuracy and a score of 30 indicating 99.9% accuracy. Table S13 presents the statistics of the clean reads.

Subsequently, we mapped clean reads to the reference genome of *Danio rerio* (assembly version GRCz11) (*59*) using Spliced Transcripts Alignment to a Reference (STAR) (*60*) with default parameters. HTSeq-count (*61*) was employed to assign the mapped reads to annotated genes and quantify the number of reads aligned to each gene, using the default parameters, with the exception of the "--stranded" option, which was set to "reverse". This configuration was necessary to differentiate between sense and antisense transcripts, as the reads were generated using a strand-specific method.

To remove unwanted technical variations introduced during sample processing, handling, or technical differences between batches, we used Surrogate Variable Analysis (SVA) from the sva (v3.54.0) R package (*62*). SVA identifies and corrects for batch effects, ensuring that statistical analyses accurately capture the true biological signal. First, we determined the optimal number of surrogate variables, which represents the best estimate of hidden confounders to be accounted for in this analysis. This was done using the num.sv function with the Buja and Eyuboglu (BE) permutation method (method="be"), applying 100 permutations (B=100) to identify the number of significant latent factors contributing to the unwanted variation. To ensure reproducibility, we set a fixed random seed (seed=100000). Next, the pre-determined optimal number of surrogate variables was used to estimate surrogate variables, representing unobserved sources of variation in the dataset. These were computed using the svaseq function with 100 iterations, which specified the number of resampling steps used to estimate the surrogate variables.

Subsequently, the estimated surrogate variables were included in downstream analyses to adjust for potential batch effects and other unmeasured confounders. To evaluate the impact of SVA correction, principal component analysis (PCA) was conducted on variance stabilizing transformation (VST) normalized expression data both before and after adjustment. PCA was performed using the prcomp function in R, and the first two principal components were visualized to assess sample clustering patterns. The resulting PCA plots illustrate the effect of SVA in reducing unwanted variation and improving clustering by biological condition (fig. S9, A and B; fig. S10, A and B; fig. S11, A and B).

For differential gene expression analysis between control and exposure conditions, we used DESeq2 (v4.3.1) in R (*63*). We first included all samples (Control, PFOS LC, PFOS HC, PFBS LC, and PFBS HC) from each generation in a single model to ensure consistent estimation of dispersion and variance across the dataset. To improve the accuracy of the differential expression analysis, both the experimental conditions and the estimated surrogate variables were included in the design formula. We then perfomed pairwise contrasts to compare control and the exposure group of interest. Genes with a Benjamini-Hochberg adjusted p-value of less than 0.05 were considered statistically significant DEGs, while, Log2 fold changes were used to quantify the magnitude of expression differences between groups. Volcano plots were generated to visualize differential expression results by plotting log2 fold change against the –log10 adjusted p-value (fig. S9, C to F; fig. S10, C to F; fig. S11, C to F). DESeq2’s robust statistical framework, advanced normalization and shrinkage estimation techniques, incorporation of independent filtering, and reliable performance compared with other methods across a variety of scenarios (*64*), make it a preferred choice for differential expression analysis in this study.

### DNA sequencing

We prepared Oxford Nanopore libraries using 5 dpf zebrafish larvae DNA from F0, F1, and F2 generations and the SKQ-NBD114.24 kit (Oxford Nanopore) according to the manufacturer’s instructions and utilizing Oxford Nanopore Barcodes provided with the kit. Two libraries were produced to accommodate all samples with at least 3 biological replicates for the same exposure and generation group and to ensure an equitable representation of different exposure groups/generations across the runs. The libraries were stored at -80°C for less than 10 days before library loading. Sequencing was performed on the PromethION platform in FLO-PRO114M (Oxford Nanopore) at SciLife lab, Uppsala, Sweden, with 20 fmol loaded per run and the flow cell loaded twice for maximizing the data output.

Next, we base-called the raw signals (pod5) generated from nanopore sequencing using the Dorado basecaller (v0.5.3) (*65*) with the super high accuracy (SUP) model dna_r10.4.1_e8.2_400bps_sup@v4.3.0_5mC_5hmC@v1, which translated the squiggle (i.e., the electric current signal) into DNA sequences. The generated reads were aligned to the reference genome of *Danio rerio* (assembly version GRCz11) (*66*) using the Dorado aligner. The aligned reads were then sorted and indexed using Samtools (*67*). We acknowledge that the sequencing depth of this dataset is limited (mean depth of 2–5× across samples) and uneven, with some genomic regions highly covered (maximum depths ranging from 1,300-7,000× across samples). Coverage exceeded 70% in all samples from F0 and F1 generations except one sample, whereas approximately half of the F2 samples showed lower coverage (43–60%). Despite variability in sequencing depth, the read coverage was higher than 80% in majority of the samples (table S5), providing adequate representation for downstream differential methylation analyses while enabling exploratory assessment of genome-wide methylation patterns. To ensure reliability and robust estimation of methylation levels, differential methylation analyses were restricted to CpG sites with a minimum coverage of three reads (3x) per group. This threshold was selected in consideration of the overall read depth of the dataset and more variable read depth inherent of long-read sequencing data generated using Oxford Nanopore Technologies (ONT) platform.

Per-site methylation calling was carried out using Modkit pileup (*68*), with the option to count only CpG motifs specified. The counts from the positive and negative strands were summed into the counts for the positive strand position, while hydroxymethylation was ignored. For the differential methylation analysis between each exposure group and their respective controls, the bedMethyl files generated from the pileup were compressed and indexed using bgzip and tabix (*69*), respectively. These files were then used as input for the Modkit DMR pair to identify DMPs, focusing only on cytosine modifications at the CpG sites. A p-value threshold of < 0.05, associated with the probability of base modification (balanced_map_p-value), was used to identify statistically significant DMPs. Subsequently, the genes associated with the identified DMPs were retrieved using the Ensembl REST API (*70*). For sites that could not be linked to any gene, the nearest transcription start site (nTSS) genes were annotated.

### Functional Enrichment Analysis

This study conducted, GSEA on Gene expression data using the clusterProfiler package (v4.14.6) (*71*) Gene ranking was performed using the statistics "stat" obtained from DESeq2, which provided a balanced combination of logFC and p-value, resulting in a single robust statistic for the GSEA. A Benjamini-Hochberg false discovery rate (FDR) threshold of 0.05 was applied to identify significantly enriched gene sets. To refine and consolidate the results, redundant Gene Ontology (GO) terms identified in the GSEA were removed using the rrvgo package (v1.14.1) (*72*) This was achieved by clustering semantically similar GO terms (*73*). Clusters were defined using similarity threshold ≥ 0.95, with lower values resulting in broader clusters. Within each cluster, GO terms were ranked by the negative log-transformed adjusted p-value from the GSEA results, and the top-ranking term with a dispensability score of 0 (range: 0–1; 0 = most representative) was retained as parent (representative) term. For the DNA methylation data, Over-Representation Analysis (ORA) was conducted using the clusterProfiler package in R (*71*).

### Correlation Analysis of DNA Methylation and Gene Expression

To assess correlations between CpG methylation and gene expression and provide insight into potential regulatory relationships between DEGs and DMG across all generations (F0, F1, F2) and exposures (PFOS, PFBS, and Control) specifically for the DEG-DMG pairs. We first integrated data from all exposure groups across generations and performed pairwise Spearman’s rank correlations between locus-level CpG methylation and their associated gene expression values across matched samples. We selected this non-parametric method based on our data distribution, which is neither normally nor linearly distributed, thus making this approach suitable for detecting monotonic relationships, particularly with modest sample sizes (n = 3 – 45 matched pairs). To stabilize the mean–variance relationship and produce a more homoscedastic distribution suitable for correlation analyses, we normalized gene expression counts using the Variance Stabilizing Transformation (VST). Pairwise Spearman’s rank correlations were then computed between locus-level CpG methylation beta values and VST-normalized expression values. We selected pairwise correlations using pairwise complete observations, to allow each CpG–gene pair to be analyzed using all available matched samples while accommodating missing data in subsets of samples.

The resulting correlation identified multiple CpG–gene pairs exhibiting statistically significant correlations at the nominal level (p<0.05). Application of Benjamini–Hochberg false discovery rate (FDR) correction substantially reduced the number of significant correlations. Nevertheless, the nominally significant correlations support a functional link between methylation and transcriptional regulation across exposure groups and generations.

To further examine the relationship between shared DEGs-DMGs across generations and exposure, we intersected the correlation results with the shared DEGs-DMGs to identify significantly correlated DEG-DMG pairs, using FDR <0.05. Only a very few DEG-DMG pairs were identified, and as such, DEG-DMG pairs with nominal P < 0.05 were retained for further exploratory analysis. Hence, we conducted over-representation analysis (ORA) using clusterprofiler in R to identify BP terms that are enriched in significantly correlated DEG-DMG pairs in F0 only. Subsequently, BP terms were reduced for redundancy using rrvgo based on semantic similarity (threshold = 0.7), selecting representative BPs according to enrichment significance. Significantly correlated DEG-DMG pairs in F1 and F2 were excluded from enrichment analysis due to insufficient gene counts (ranging from one in the F1 groups to a maximum of 15 in an F2 exposure group) for reliable statistical testing.

### Association Analysis Between F2 Behavior and F0/F1/F2 DNA Methylation

To associate behavioral outcomes in the unexposed generation (F2) with specific CpG sites, we first used Modkit pileup (*68*) to estimate per-site methylation counts for each sample, integrated counts from all exposure conditions for each generation, and processed to obtain site-specific methylation proportions (β-values), defined as the ratio of methylated counts to total counts for each CpG site. Later, we filtered CpG sites to retain only those with a minimum coverage of ≥5 reads in at least 50% of samples. To avoid degenerate values, we then identified and excluded CpG sites with zero variance across samples from each generation.

To satisfy the assumptions of beta regression, beta values equal to 0 or 1 were transformed to fall within the open interval (0,1) using the “squeezing” transformation 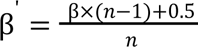 (*74*), where β’, β and *n* are transformed beta-values, raw beta-values and the number of samples, respectively. M-values were then computed as 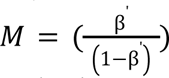 for surrogate variable estimation. Potential unobserved confounders were estimated using surrogate variable analysis (SVA) in the R package sva (*62*). Due to the limited sample size, only one surrogate variable (SV1) was retained to prevent overfitting and preserve biological signals. Collinearity between SV1 and known batch effects was assessed using Pearson correlation and one-way ANOVA (*aov*(*sv*∼*Batch*)). Known batch effect was included as a covariate only if the correlation with SV1 was <0.2 and the ANOVA *p*-value was ≥0.01.

Subsequently, we estimated associations between methylation and behavioral outcomes using beta regression implemented in the R package betareg (v3.2.4) (*75*) to model methylation proportions (β′) for each generation as a function of F2 behavioral outcomes and covariates (i.e SV1 and known batch effect) for each generation. The model was designed as follows: *Model*_*x*_ <− *betareg*_(_β’_*x*_ ∼ *F*2_*behavior* + *SV*1_*x*_ + *Batch*_*x*)_. Where x is the generation (F0, F1, F2). The unadjusted p-value <0.05 was used to select statistically significant CpG sites referred to as BAPs. Although multiple-testing correction was explored, very few CpG sites remained significant after adjustment, mostly in F2 association analysis; therefore, unadjusted p-values were reported to capture potential biological signals.

### Identification of F0 Epigenetic Signatures Associated with Transgenerational Outcomes

To identify epigenetic signatures in the F0 generation that could predict behavioral outcomes in subsequent generations, we first identified overlapping sites between the BAPs detected using Models F0, F1, and F2. This step identified common and specific changes across generations. Next, we defined the epigenetic signature of transgenerational epigenetic inheritance (TEI) as the set of BAPs commonly identified across all models. Each BAP was mapped to its corresponding gene using the Ensembl REST API (*70*). For BAPs located in the intergenic region, genes with the nearest transcription start site (TSS) were assigned. Genes that lacked an HGNC-approved gene symbol were retained and reported using their Ensembl gene identifiers. To evaluate the functional relevance of the shared BAPs, we performed over-representation enrichment analysis using the clusterProfiler package (*71*). Enriched functional categories were subsequently summarized using the rrvgo package (*72*), which reduces redundancy among enriched terms.

### Statistical analysis

All statistical analyses were conducted in R version 4.4.1 (14 June 2024) using publicly available packages. Unless otherwise indicated, p-values derived from statistical analyses were corrected for multiple hypothesis testing using the Benjamini–Hochberg false discovery rate method (*76*), and adjusted p-values were used to determine statistical significance. For all analyses, we defined statistical significance as adjusted p-value < 0.05, except where noted.

## Supporting information

https://docs.google.com/document/d/1acW8eUcjEXLVV2zv_S2ijh05QVrzhkBi/edit?usp=sharing&ouid=115672375919768063843&rtpof=true&sd=true

## Acknowledgments

We thank Emmanouil Tsakoumis (PhD) for his precious help handling zebrafish and collecting samples.

## Ethics statement

All experimental procedures performed on zebrafish larvae and juveniles were in accordance with national and international laws, regulations, and agreements. Zebrafish husbandry and breeding were conducted under permits issued by the Swedish Board of Agriculture, Jönköping, Sweden (# 5.2.18-12628/17, # 5.2.18-12630/17) and ethical approval from Linköping’s animal test ethics committee (ID 1166).

## Author contributions

Conceptualization: AZO, MD, MF, PA, JR, NS and SHK

Methodology and Investigation: AZO, MD and MF

Data curation: AZO, MD and MF

Investigation and Formal analysis: AZO, MD, MF, JZ and CY

Software: AZO

Visualization: AZO and MF

Supervision: MF, JR, PA and SHK

Writing—original draft: AZO, MD and MF

Writing—review & editing: all authors

Project administration, Funding acquisition and Resources: JR and SHK

## Data and materials availability

All data required to evaluate the conclusions of this study are included in the main text and/or the supplementary materials. All raw data and laboratory materials used and generated in this study will be deposited in a publicly accessible database upon publication. Analysis scripts, including code used for data processing, data analysis and figure generation, are available on GitHub at the following link: https://github.com/adeolu1/Differential_Gene_Expression_Analysis.

