## Supplementary material for "Epigenetic changes induced by developmental PFAS exposure in zebrafish associate with behavioral alterations in unexposed offspring": https://docs.google.com/document/d/1acW8eUcjEXLVV2zv_S2ijh05QVrzhkBi/edit?usp=sharing&ouid=115672375919768063843&rtpof=true&sd=true: Preprint_supplementary_materials.docx.pdf

Adeolu Z. Ogunleye *et al.*

**This PDF file includes:**

Figs. S1 to S11
Table S9B

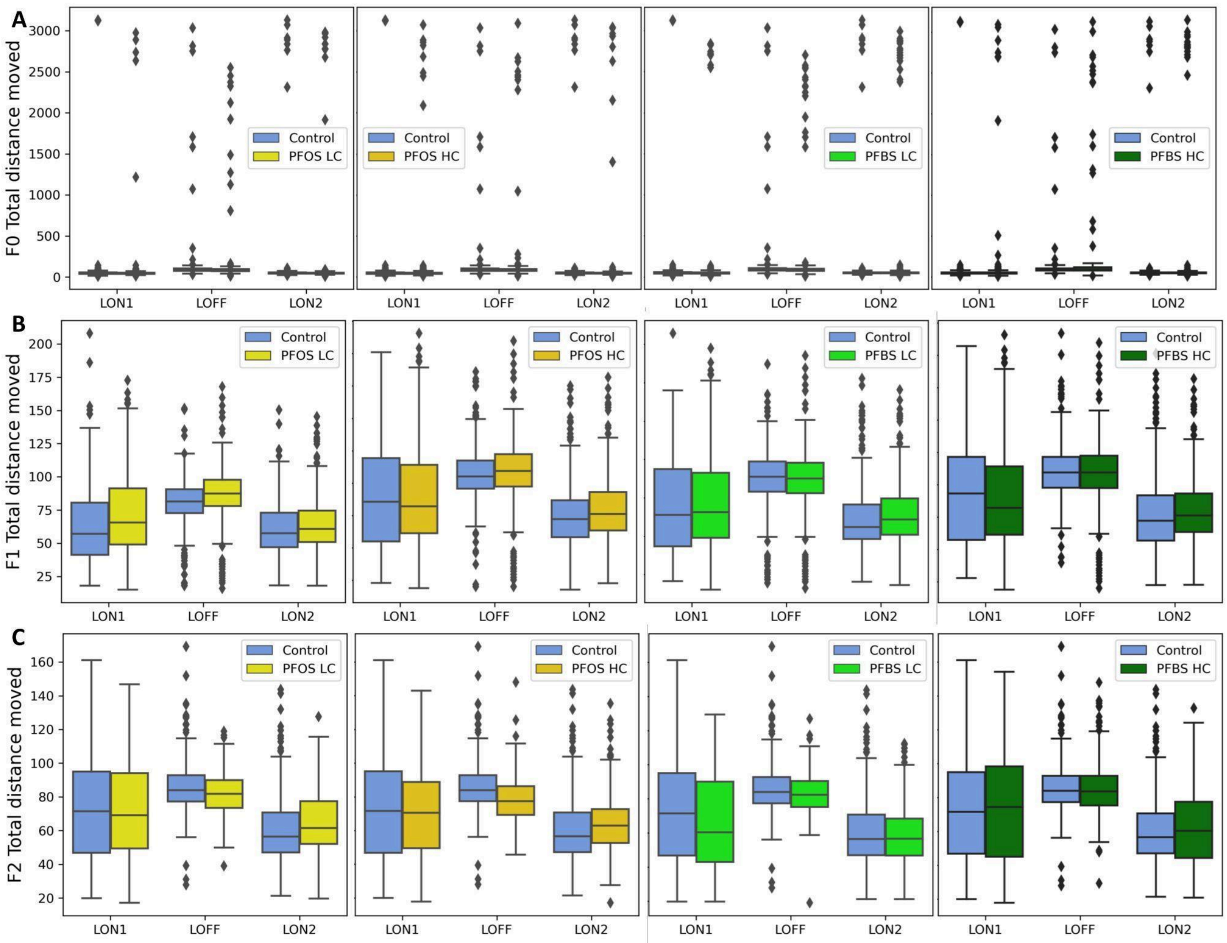

**Fig. S1: Behavioral analysis in the F0, F1, and F2.**

Total distance moved by larvae at 5 dpf during the LMPR test and the phases LON1 (acclimatization); LOFF (dark stimulus); LON2 (recovery) for all exposure conditions: Control (blue), PFOS LC (yellow), PFOS HC (orange), PFBS LC (light green), and PFBS HC (dark green), and are displayed for the F0 (A), F1(B), and F2 (C) generations (top, middle, and bottom panel respectively).

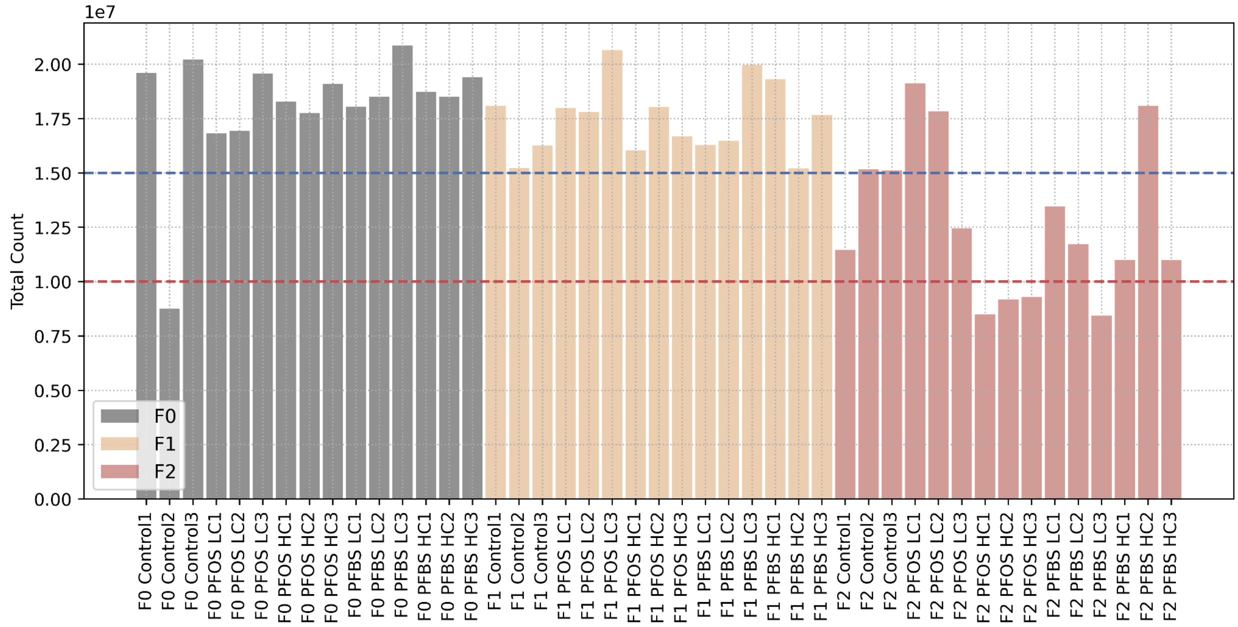


**Fig. S2: Total CpG sites detected per sample by Oxford Nanopore Technologies (ONT) sequencing.**
Barplot showing the total number of sequencing reads generated per sample by ONT sequencing. Each bar represents an individual sample (n = 45), with colors indicating generations. The plot highlights variability in the total number of CpG sites identified across samples, providing an overview of methylation coverage across generations.

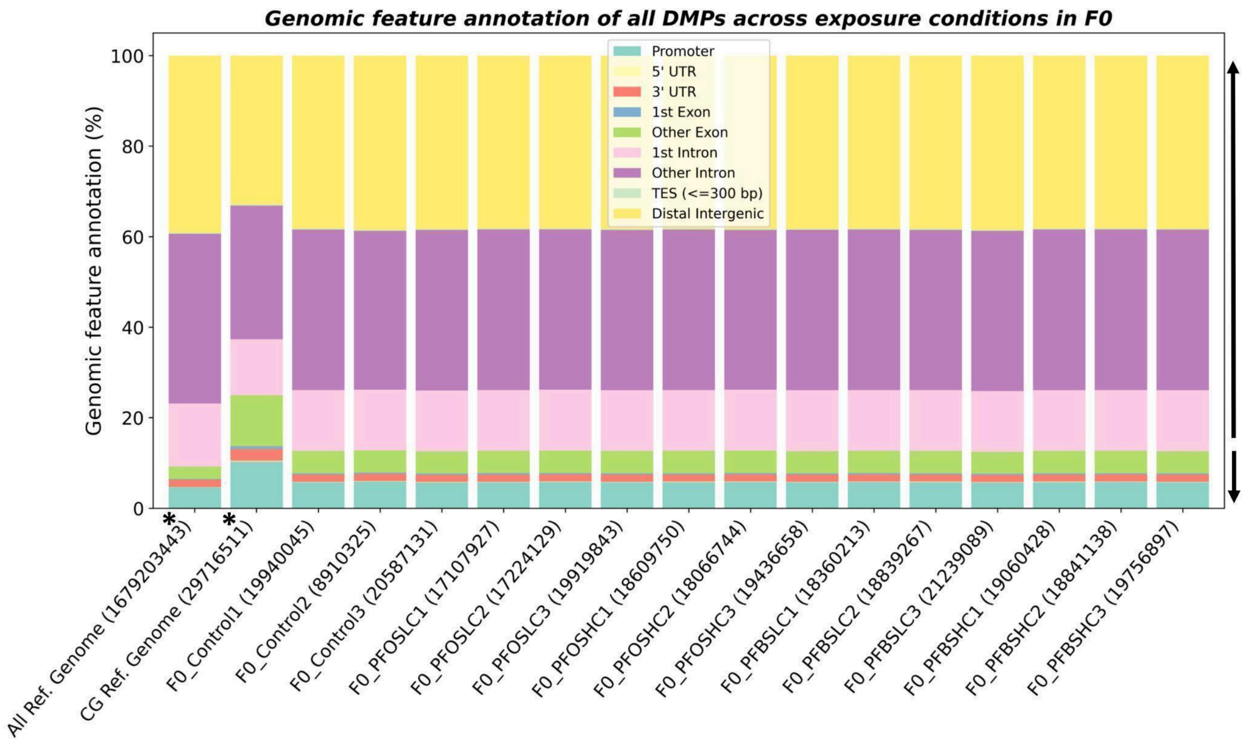
Genomic feature annotation of all DMPs across exposure conditions in F0
Genomic feature annotation (%)
Promoter
5' UTR
3' UTR
1st Exon
Other Exon
1st Intron
Other Intron
TES (<=300 bp)
Distal Intergenic
All Ref. Genome (1679203443)
CG Ref. Genome (29716511)
F0_Control1 (19940045)
F0_Control2 (8910325)
F0_Control3 (20587131)
F0_PFOSLC1 (17107927)
F0_PFOSLC2 (17224129)
F0_PFOSLC3 (19919843)
F0_PFOSHC1 (18609750)
F0_PFOSHC2 (18066744)
F0_PFOSHC3 (19436658)
F0_PFBSLC1 (18360213)
F0_PFBSLC2 (18839267)
F0_PFBSLC3 (21239089)
F0_PFBSHC1 (19060428)
F0_PFBSHC2 (18841138)
F0_PFBSHC3 (19756897)

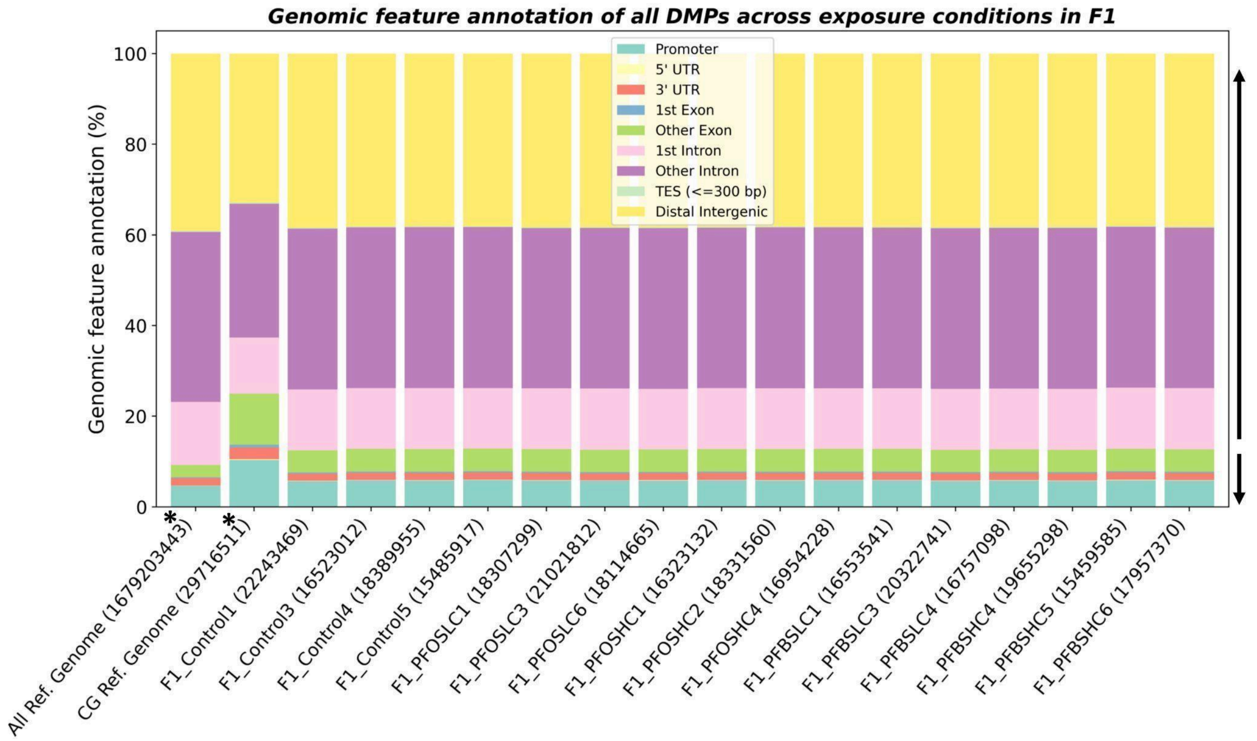
Genomic feature annotation of all DMPs across exposure conditions in F1
Genomic feature annotation (%)
Promoter
5' UTR
3' UTR
1st Exon
Other Exon
1st Intron
Other Intron
TES (<=300 bp)
Distal Intergenic
All Ref. Genome (1679203443)
CG Ref. Genome (29716511)
F1_Control1 (22243469)
F1_Control3 (16523012)
F1_Control4 (18389955)
F1_Control5 (15485917)
F1_PFOSLC1 (18307299)
F1_PFOSLC3 (21021812)
F1_PFOSLC6 (18114665)
F1_PFOSHC1 (16323132)
F1_PFOSHC2 (18331560)
F1_PFOSHC4 (16954228)
F1_PFBSLC1 (16553541)
F1_PFBSLC3 (20322741)
F1_PFBSLC4 (16757098)
F1_PFBSHC4 (19655298)
F1_PFBSHC5 (15459585)
F1_PFBSHC6 (17957370)

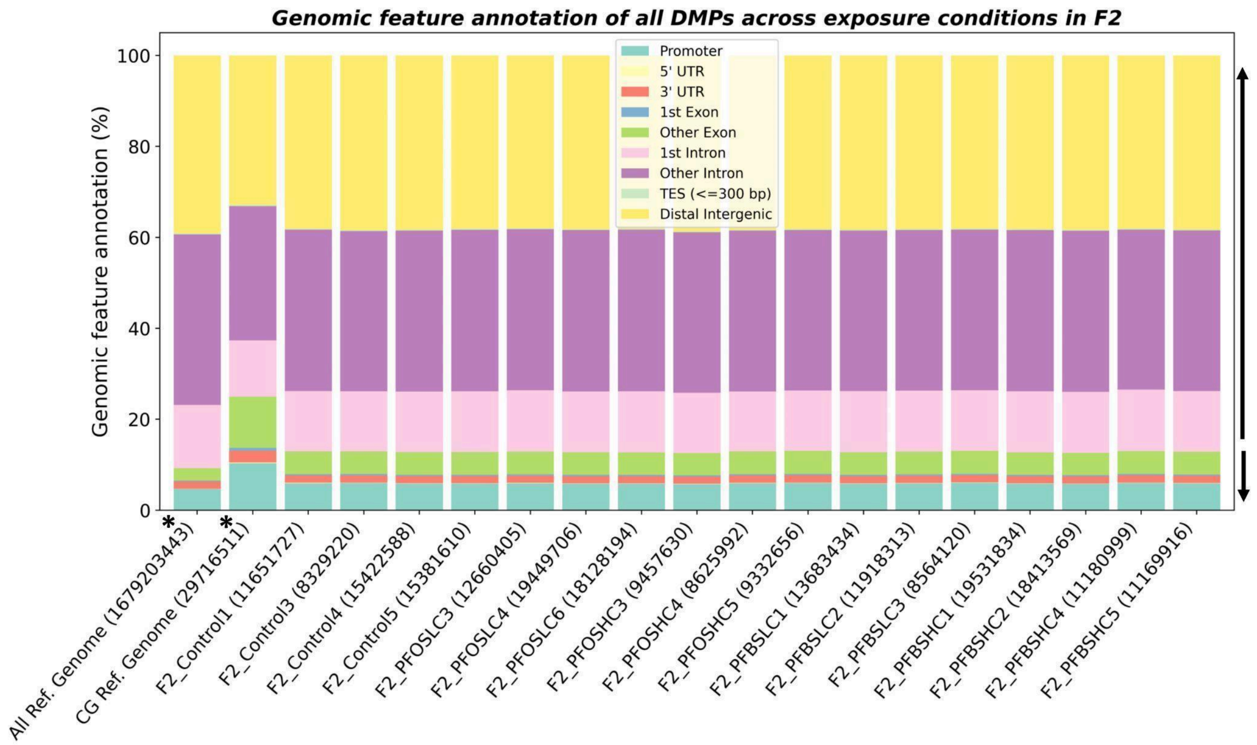


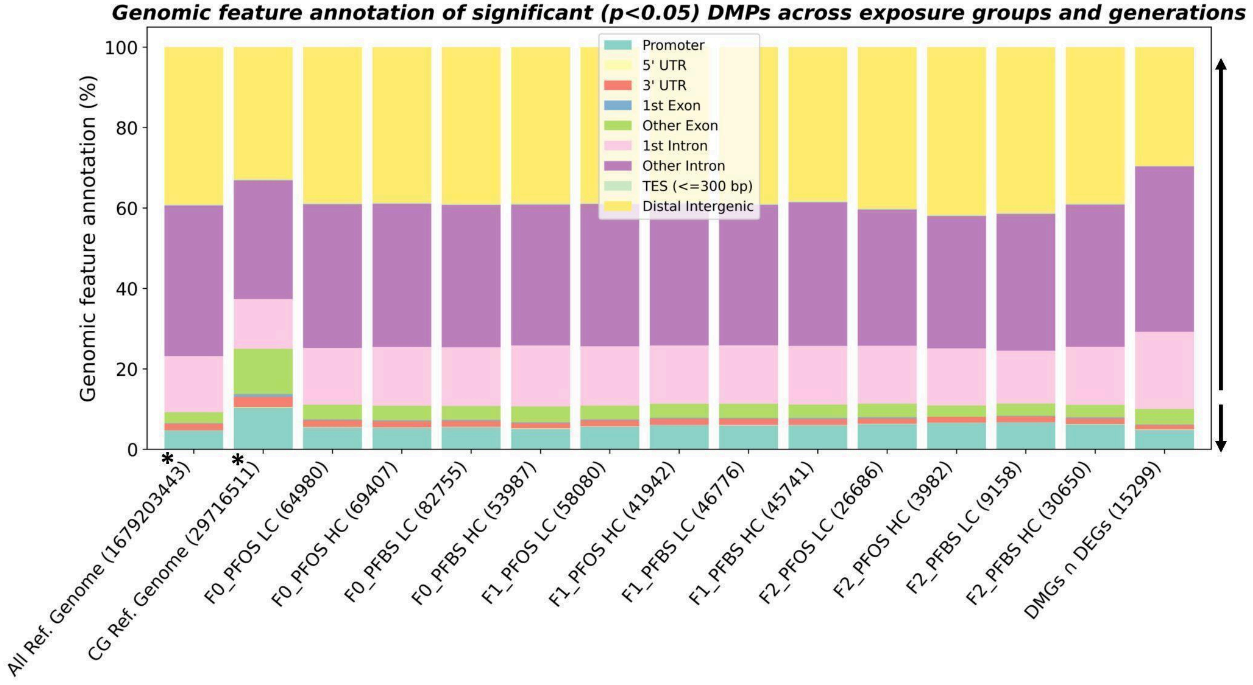


**Fig. S3: Genomic feature annotation.**

Percentage (%) of bases mapped to specific components of the Zebrafish reference genome among all samples across exposure conditions in F0, F1, F2, and significance ($P < 0.05$) differentially methylated positions (DMPs), respectively. Promoter was defined as −1000 to +500 bp relative to the transcription start site (TSS). All Ref. Genome* represents all nucleotide bases (A, T, C, and G) in the Zebrafish reference genome and CG Ref. Genome* represents the cytosine–guanine (CG) bases in the CpG context.

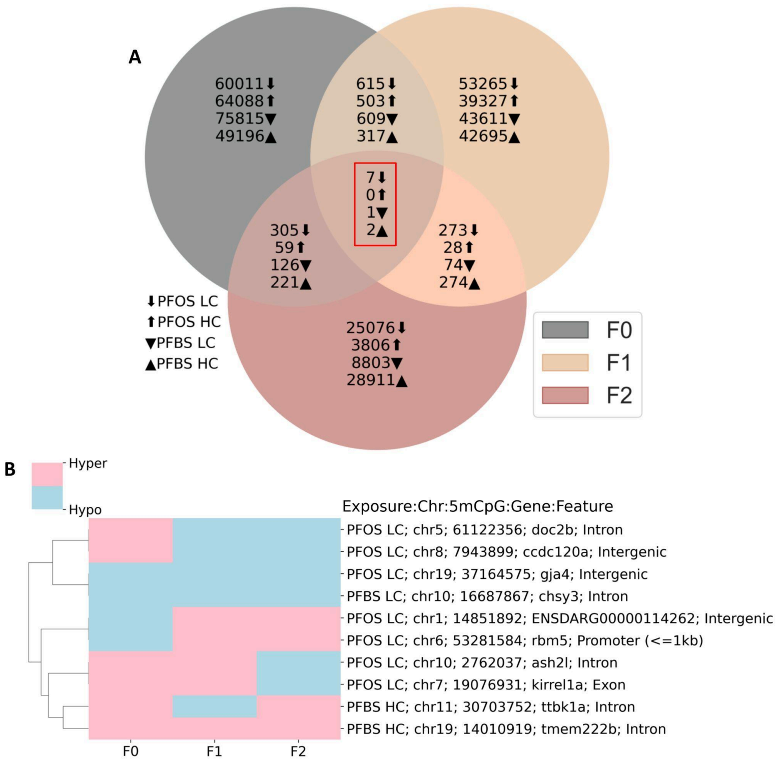


**Fig. S4. Overlap of DMPs across generations.**
(A) Venn diagram showing the number of shared and unique DMPs identified in each generation. The intersection represents DMPs common to all generations. (B) Clustering heatmap of the overlapping DMPs, corresponding to the regions highlighted in the red square in A. Rows represent DMPs, and columns represent generations, with pink and blue indicating hypermethylation (hyper) and hypomethylation (hypo), respectively.

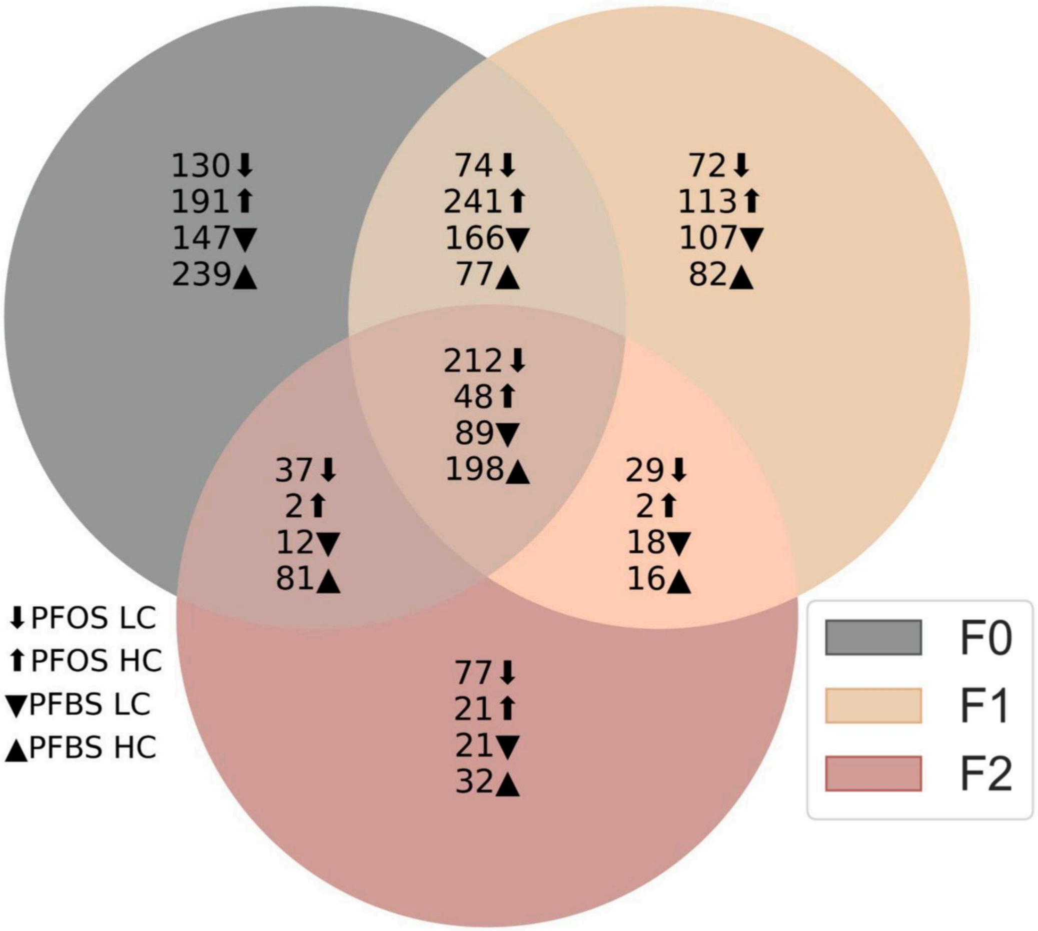


**Fig. S5: Number of biological processes (BP) enriched in differentially methylated genes (DMGs) across generations and exposure conditions.**
Venn diagram showing the number of significantly (FDR < 0.05) enriched BPs in DMGs of PFOS LC, PFOS HC, PFBS LC, and PFBS HC conditions across F0, F1, and F2 generations.

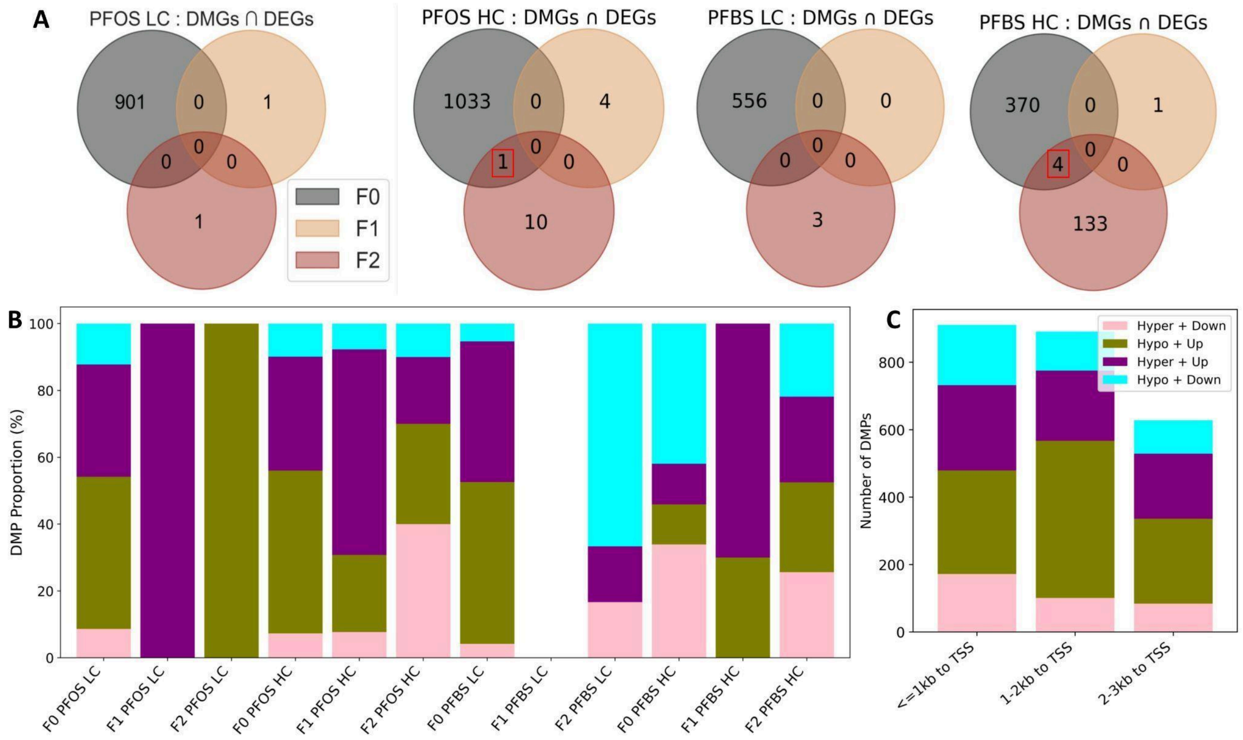


**Fig. S6: Distribution and genomic features of shared DEGs and DMGs.**

(A) Overlap between DEGs and DMGs across exposure groups and generations. The red square highlights the number of DEGs-DMGs that overlap transgenerationally between F0 and F2. (B) Percentage distribution of total DMPs across genomic features, showing the proportion of DMPs occurring in each feature relative to the total number of DMPs. (C) Distribution of Methylation–Expression patterns in promoter DMPs to the transcription start site (TSS) upstream and downstream.

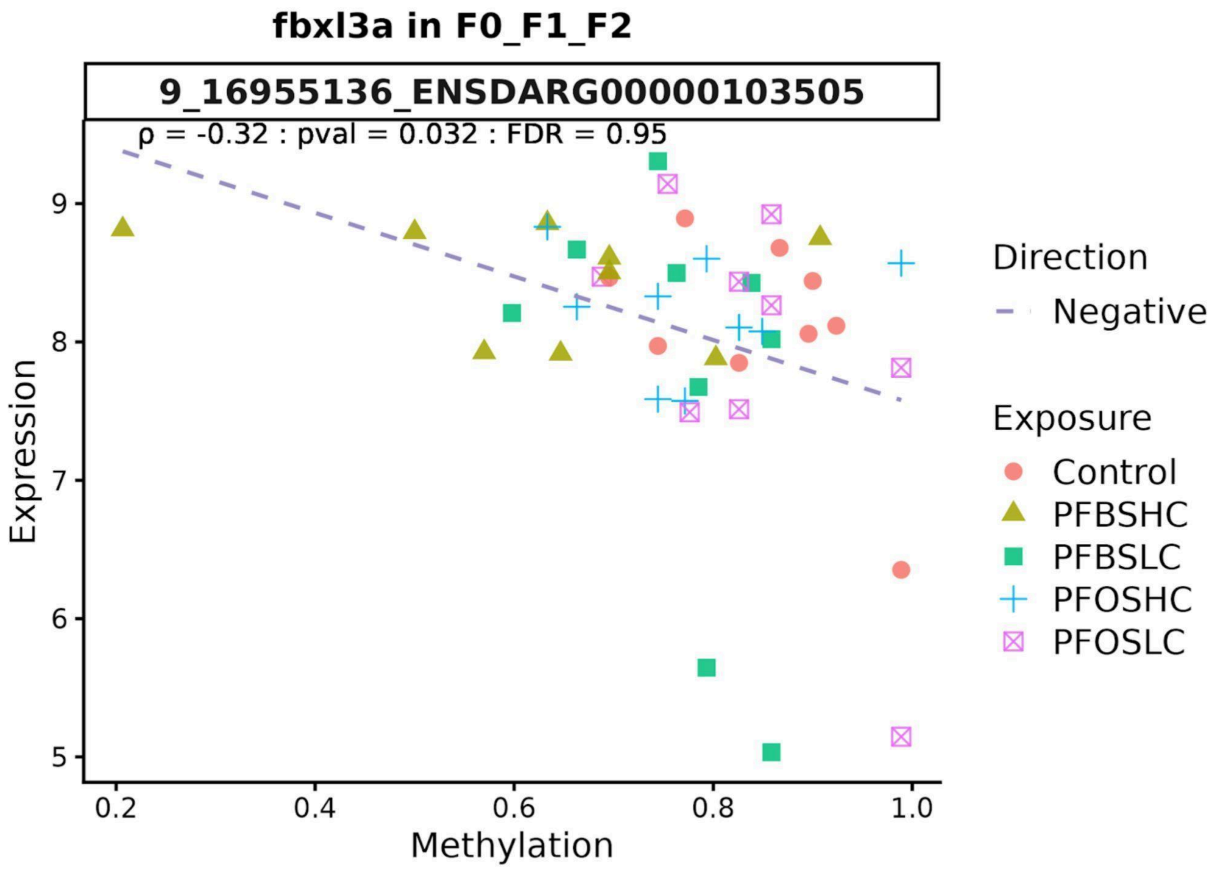


**Fig. S7: Inverse correlation between gene expression and DNA methylation at the fbxl3a locus.**
Expression of the fbxl3a inversely correlated with DNA methylation levels across samples. The scatter plot shows normalized fbxl3a expression plotted against corresponding methylation levels at CpG sites annotated to fbxl3a. Each point represents an individual sample. A linear regression model (dotted line). Statistical analysis reveals a significant negative correlation (Spearman's correlation coefficient = 0.32, p = 0.032).

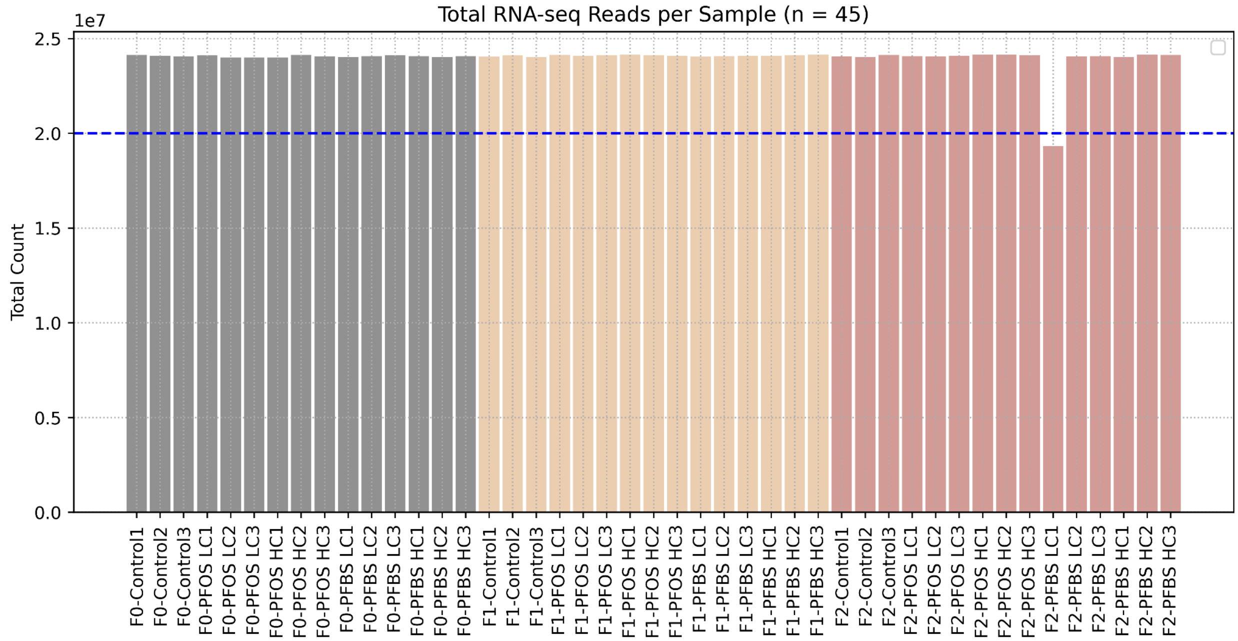


**Fig. S8: Sequencing depth across samples.**
Barplot presenting the total number of sequencing reads generated per sample. Each bar represents an individual sample, colored by generation. On average, approximately 24 million paired-end reads were obtained per sample, indicating consistent sequencing depth across all samples.

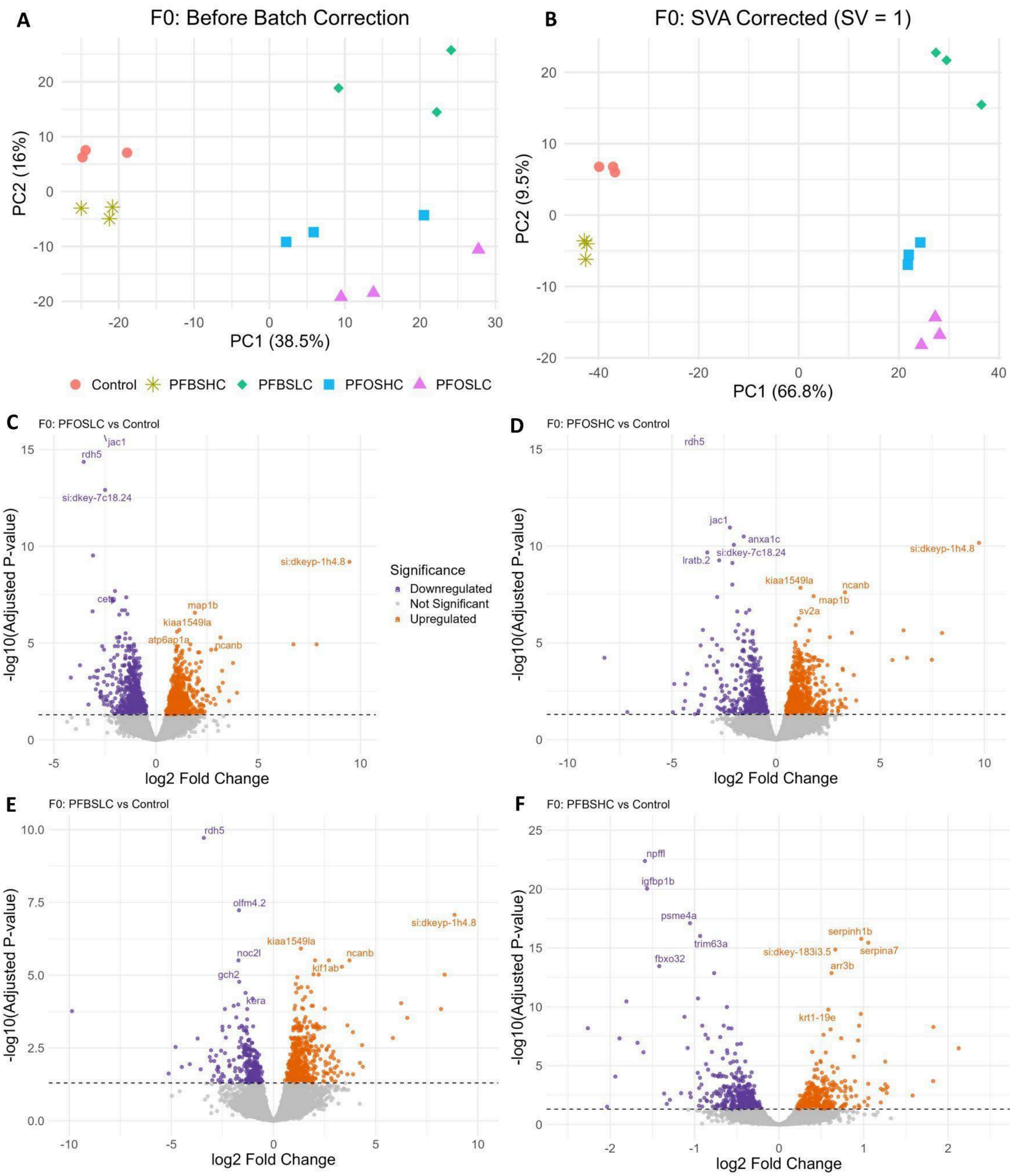


**Fig. S9: Principal component analysis and differential expression across exposure groups in F0.**
(A and B) Principal component analysis (PCA) of normalised F0 gene expression data was performed (A) before and (B) after surrogate variable analysis (SVA). Each point represents an individual sample, colored and shaped by exposure groups. SVA correction reduces variability attributable to latent confounding factors and improves clustering according to exposure condition. In the F0 data, one surrogate variable (SV1) was identified as sufficient to capture hidden sources of variation. This SV was subsequently included in the differential expression model to adjust for latent sources of variation. The inclusion of this SV improved sample clustering as observed in the PCA plots. (C to F) Volcano plots showing differential gene expression results for pairwise comparisons between control and each exposure group (PFOS LC, PFOS HC, PFBS LC, and PFBS HC), as determined by DESeq2. Log2 fold change is plotted against the –log10 adjusted p-value (false discovery rate, FDR). Each point represents a gene. Genes meeting the significance threshold (FDR < 0.05) are highlighted. Significantly downregulated and upregulated genes are shown in purple and orange, respectively, while non-significant genes are shown in grey in each comparison. The top five upregulated and top five downregulated genes, ranked by statistical significance, are labelled.

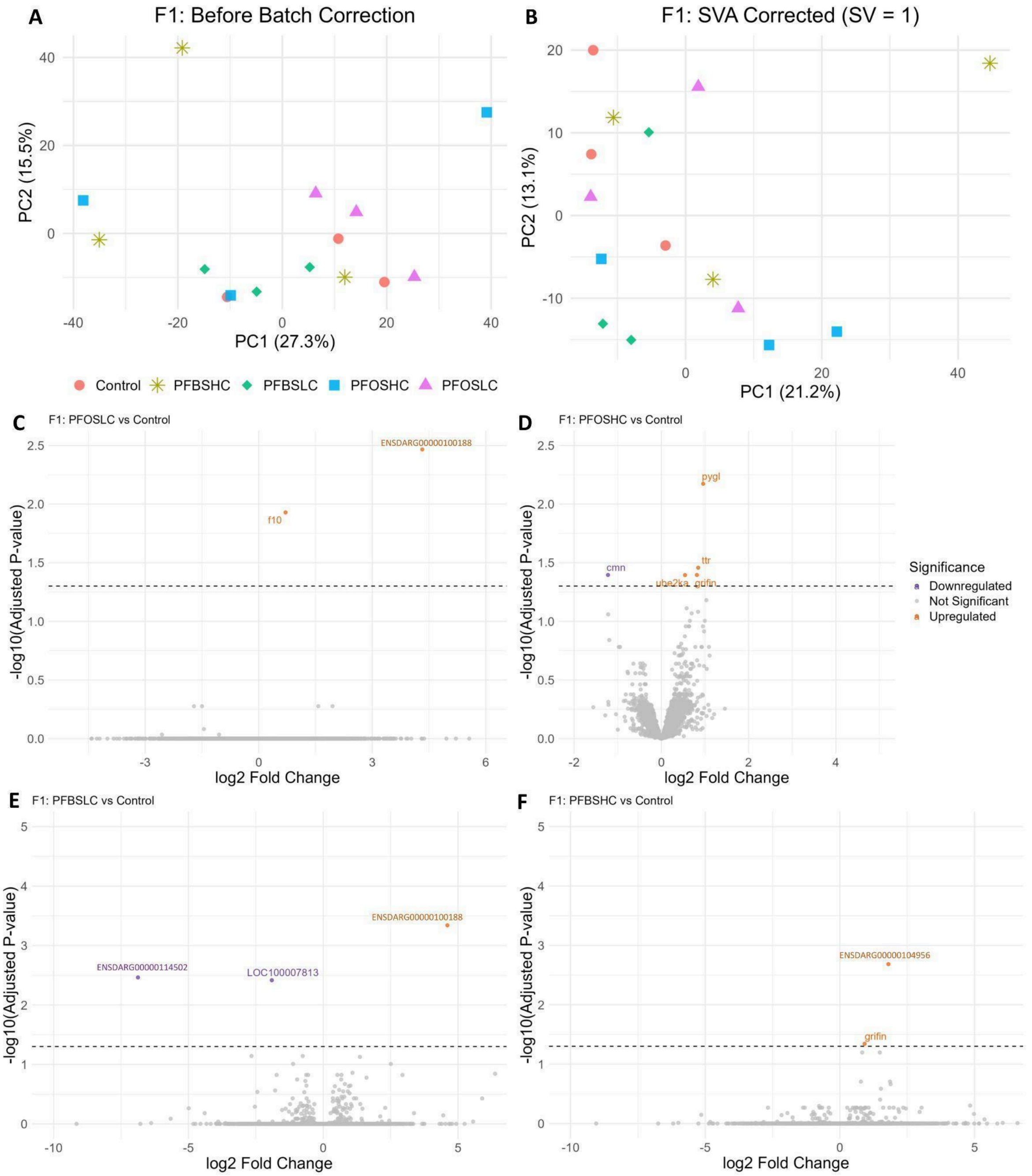


**Fig. S10: Principal component analysis and differential expression across exposure groups in F1.** (A and B) Principal component analysis (PCA) of normalised F1 gene expression data was performed (A) before and (B) after surrogate variable analysis (SVA). Each point represents an individual sample, colored and shaped by exposure groups. SVA correction reduces variability attributable to latent confounding factors and improves clustering according to exposure condition. For F1 gene expression data, one surrogate variable (SV1) was identified as optimal and was incorporated into downstream differential expression analyses to correct for latent confounding factors. The inclusion of this SV improved sample clustering as observed in the PCA plots. (C to F) Volcano plots showing differential gene expression results for pairwise comparisons between control and each exposure group (PFOS LC, PFOS HC, PFBS LC, and PFBS HC), as determined by DESeq2. Log2 fold change is plotted against the –log10 adjusted p-value. Each point represents a gene. Genes meeting the significance threshold (adjusted p-value < 0.05) are highlighted. Significantly downregulated and upregulated genes are shown in purple and orange, respectively, while non-significant genes are shown in grey in each comparison. All significantly upregulated and downregulated genes, ranked by adjusted p-value, are labelled.

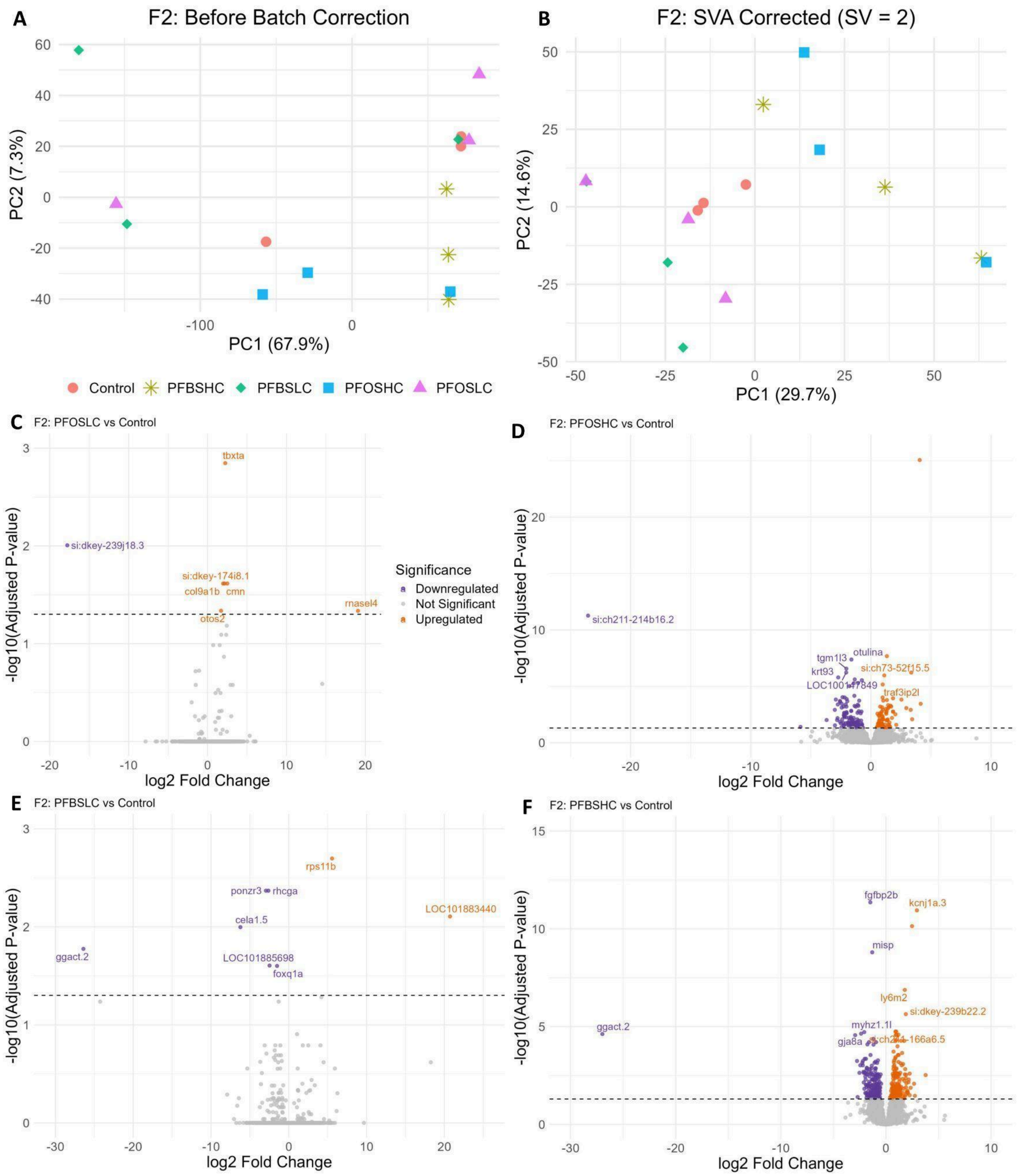


**Fig. S11: Principal component analysis and differential expression across exposure groups in F2.**
(A and B) Principal component analysis (PCA) of normalised F2 gene expression data was performed (A) before and (B) after surrogate variable analysis (SVA). Each point represents an individual sample, colored and shaped by exposure groups. SVA correction reduces variability attributable to latent confounding factors and improves clustering according to exposure condition. Based on F2 gene expression data, two surrogate variables (SV1 and SV2) were identified as optimal and were included in downstream modelling to correct for latent confounding factors. The inclusion of these SVs improved sample clustering as observed in the PCA plots. (C to F) Volcano plots showing differential gene expression results for pairwise comparisons between control and each exposure group (PFOS LC, PFOS HC, PFBS LC, and PFBS HC), as determined by DESeq2. Log2 fold change is plotted against the –log10 adjusted p-value. Each point represents a gene. Genes meeting the significance threshold (adjusted p-value < 0.05) are highlighted. Significantly downregulated and upregulated genes are shown in purple and orange, respectively, while non-significant genes are shown in grey in each comparison. The top five upregulated and top five downregulated genes, ranked by adjusted p-value, are labelled to highlight the most significant expression changes; where fewer than five genes met the significance threshold, all such genes were labelled.

**Table S9B: Summary of DNA methylation and gene expression correlation analysis**

| Generation | Exposure | # Samples | # Total Samples | # CpG sites | # Genes |
|---|---|---|---|---|---|
| F0 | Control<br>PFOS LC<br>PFOS HC<br>PFBS LC<br>PFBS HC | 15 samples<br>(3 per group) | 45 | 23712971 | 32500 |
| F1 | | 15 samples<br>(3 per group) | | | |
| F2 | | 15 samples<br>(3 per group) | | | |

***Summary of the statistically significant results (P<0.05 vs FDR <0.05)***

| **% Missing samples** | **# Samples (>=)** | **# Correlated CpGs pval<0.05** | **# Correlated Genes pval<0.05** | # Correlated CpGs FDR<0.05 | # Correlated Genes FDR<0.05 |
|---|---|---|---|---|---|
| All Significant | 3 | 946801 | 27756 | 7060 | 2145 |
| No missing sample | 45 | 1922 | 286 | 1 | 1 |
| 5% missing | 42 | 6317 | 1072 | 5 | 5 |
| 10% missing | 40 | 17071 | 4188 | 40 | 35 |
| 15% missing | 38 | 99131 | 16152 | 356 | 228 |
| 25% missing | 33 | 521949 | 25826 | 2149 | 840 |
| 50% missing | 22 | 894235 | 27463 | 3751 | 1346 |

**Pairwise correlation analysis between locus-level CpG methylation and gene expression.** The table summarises the proportion and number of samples with pairwise observations used for correlation analysis (minimum n = 3). "No missing sample" indicates that all 45 samples had pairwise values, whereas "5% missing" indicates that at least 42 of 45 samples had pairwise values and were included in the analysis. "#Samples (≥)" denotes the minimum number of samples with pairwise observations used for correlation.
